# Category learning disentangles representation of trial events in hippocampus CA1

**DOI:** 10.64898/2025.12.01.690977

**Authors:** Laura Sainz Villalba, Ioana Calangiu, Roman Boehringer, Valerio Mante, Benjamin F. Grewe

## Abstract

The hippocampus (HC) is known to encode task-relevant variables, including latent ones, capturing and parsing critical information from sequences into episodes. This is thought to be the basis to form a cognitive map of possible abstract states beyond mere perceptual details, akin a state machine, with predictive value, contributing to an internal world model. However, its specific role in categorization tasks in general, where learning a category may depend on independent, unordered (non-sequential) experienced examples, remains unclear.

Here, we investigate CA1 population coding during a categorization paradigm in mice in combination with calcium imaging at different stages of category training with interleaved generalization tests.

Our results reveal that hippocampal coding changes critically throughout the different stages of categorization training. Specifically, the disentangling of choice and outcome variables emerges as factorized, abstract, representations and fundamentally distinguishes simple discrimination from categorization training. Furthermore, this factorized geometry relates to improved behavioral performance. Our findings suggest that trial encoding on HC adapts in response to the category structure presented by representing critical events into the appropriate computational format to support generalization of category membership.

## Main

Animals and humans rely on past experience to generalize to novel situations [1]. Hippocampus (HC) is thought to support this ability by encoding relevant events, building what is known as a cognitive map [2] in what is described as a supporting substrate for schemas, concepts and categories. Cognitive maps are proposed as neural representations that support flexible behavior, enabling fast inference from sparse observations [2,3]. While originally linked to spatial navigation alone, this framework has expanded to non-spatial cognition: recent work has demonstrated that non-spatial variables such as sound frequency [4], reward [5], and accumulated evidence [6] are also encoded in the HC if relevant to the task. A common feature of the encoded variables in these experiments, including navigation tasks, is that they present an informative sequential structure in the task paradigm [7,8]. Such sequential regularities supply the HC with a scaffold for organizing experiences into states, thereby facilitating efficient computation. For example, in Aronov et al [5], rodents had to press a lever at a specific tone frequency to receive reward. Tone frequencies were presented as stereotypical ordered frequency swipes. This fixed transition structure effectively allowed treating frequency as a spatial-like variable, enabling the HC to anchor reward events to “locations” along this tone frequency dimension.

These findings suggest that cognitive maps act as a general coding mechanism, covering any domain that shows similar properties to physical space [9]. Indeed, broader understanding describes the fundamental function of the hippocampus as to encode and predict sequential order of experiences [10] capturing the abstract structure of tasks into states, separated from specific sensory information, enabling generalization [11].

In this setting, hippocampal activity then organizes experiences into ordered abstract trajectories, described through successor-like representations that capture the dependency structure of events as a predictive map.

However, we are still yet to fully characterize hippocampal role in categorization tasks (in general with no sequential structure), where animals must learn to infer category membership (an abstract variable) from previous examples that are experienced as independent, unordered, learning episodes of category structure. In these tasks, there is no transition structure to exploit [9] since neither the stimulus dimensions nor their categorical labels follow a sequence. Key evidence about the nature of hippocampal representations serving as a basis for category learning, remains elusive. Specifically, does CA1 activity represent event states in line with its described role as a state machine even in the absence of sequential structure on category-relevant dimensions? Does hippocampal population activity reflect changes in category structure, even capturing similarity according to category membership?

Here we address these issues by performing large-scale neuronal recordings in Hippocampus CA1 during a categorization task in mice. Concretely, we used a binary auditory low-vs-high tone categorization task defined on a single continuous axis (tone frequency) experienced in no particular sequence across trials. This removes transition regularities to avoid providing a reliable sequential structure that could be leveraged for tone frequency relating to category. We then staged the learning explicitly so that learning of category structure would be separated from learning the trial structure. In the first stage (auditory discrimination training), mice were presented only with one stimulus per category, such that performance would depend on learning trial structure and discriminating between the two stimuli, and not necessarily on understanding of category structure. As the paradigm progressed to categorization training, we included more stimuli items per category and improving performance was now really dependent on learning the category structure. Additionally, we explicitly checked category understanding with interleaved tests, to uncover any generalization abilities induced from training stages (especially after simple discrimination training).

Collectively, our results suggest that representation of variables (like choice and outcome) changes with the complexity of the category structure presented and not purely according to trial structure dependencies of events. Our findings provide insight into potential neuronal mechanisms in the hippocampus to provide episodic memory in a suitable representation format at the population level that would allow other areas to extract critical information of experienced events fostering category learning.

## Results

To study the role of Hippocampus in learning a category structure we developed a novel staged paradigm for mice based on a one-dimensional auditory binary categorization task with an absolute single boundary. Each full-training session consisted of about 100 trials where on each trial a brief tone stimulus was presented and the mouse subject reported the category (A or B) assigned to the tone by licking either to the left or to the right spout after a delay period. Auditory stimuli consisted of a discrete set of pure tones ranging from 2000Hz to 20000Hz in frequency with a power-law spacing (See Methods). The underlying ground truth for correct categorization was based on a linear split between low and high frequencies (**Fig. 1A**). When the reported choice was correct, mice received a water drop as reward, whereas for the incorrect responses a timeout followed (**Fig. 1B**, Supplementary Movie 1).

**Figure 1:**
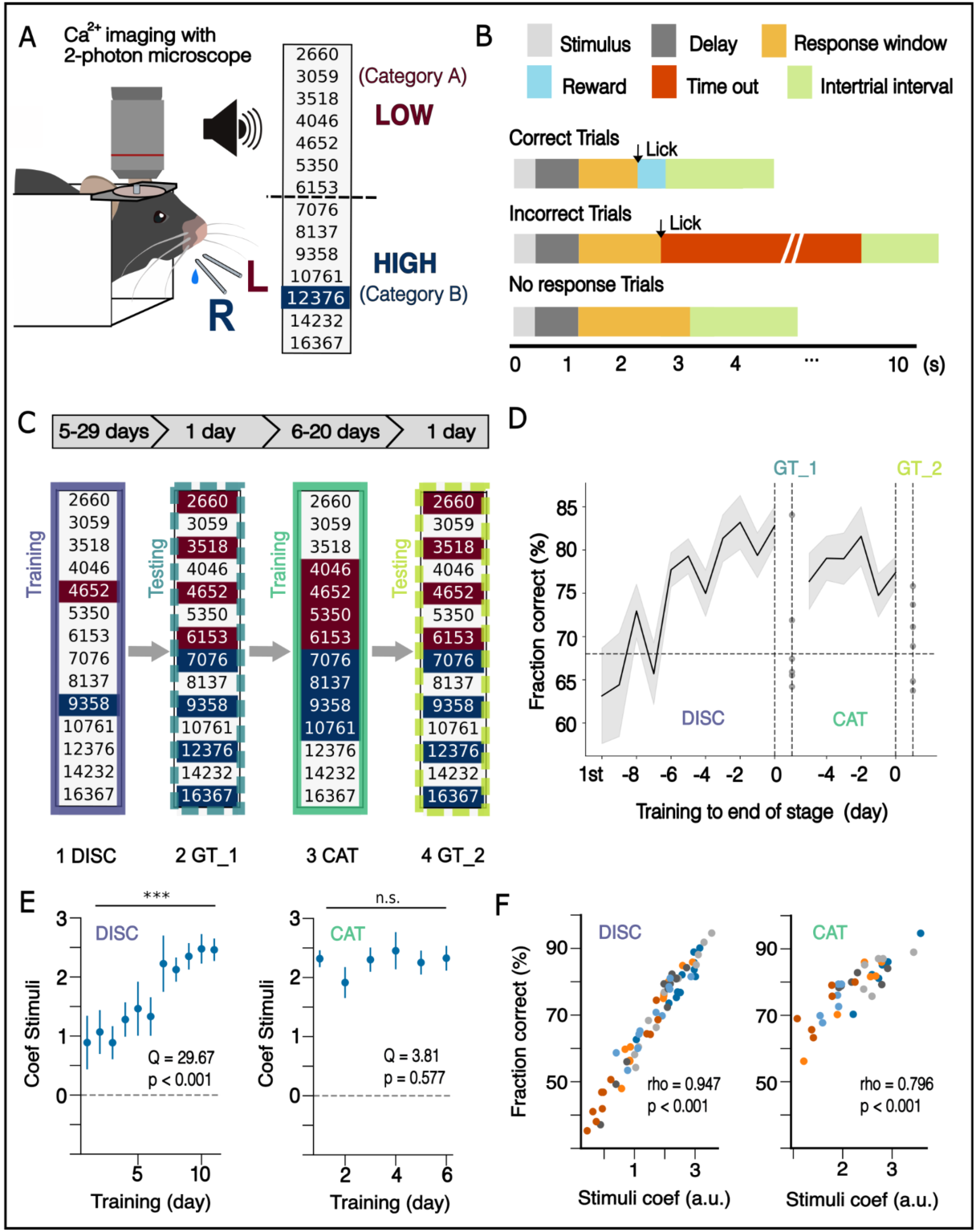
Task paradigm, stages and overview of learning. (A) Task paradigm consisting of a one-dimensional auditory binary categorization upon tone frequency between low and high tones. (B) Trial structure. Each trial begins with a tone presentation preceded by a jittered pre-stimulus interval. A delay period follows until the water ports become available during a response window. Depending on the choice and stimulus, a water reward or timeout is delivered (See Methods). (C) Different stages of training. Stimuli available at each stage are highlighted. Discrimination (1-DISC) and categorization (3-CAT) correspond to training stages, whereas (2-GT_1) and (4-GT_2) correspond to generalization tests performed only for one session. (D) Cohort learning curve for fraction of correct throughout stages of the paradigm aligned to the last day of each stage. In the discrimination stage, the first day is plotted as a reference as 1st. (E) Trend of coefficient for stimuli regressor in logistic regression for choices (mean +- standard deviation) tested in discrimination (left) showing upward trend (n=6, p < 0.001, Friedman rank sum test) and categorization stage (n=6, p = 0.577, Friedman rank sum test) showing stability (right). (*: p < 0.05, **: p < 0.01, ***: p < 0.001) (F) Correlation scatter plot of fraction of correct vs stimuli regressor coefficient for each session color coded according to subject id. For both training stages (discrimination and categorization), the Spearman rank coefficient was high and significant.

The staging consisted of two learning/training stages and two testing ones (generalization tests, unique sessions). The first training stage, *discrimination learning*, comprised a pair of stimuli, one per category, to make mice get familiar to the task trial structure. To speed up learning, we chose two tone frequencies *easy* to distinguish. With a mean training time of 16.3 +- 2.4 days, mice reached 81.1% +- 2.4% fraction of correct choices, and 1.9 +- 0.2 for discrimination d’ (mean +- s.e.m, **Fig. 1C, 1D, S1**) (see Methods). After being proficient on this stage, mice advanced to the first testing stage, the *generalization test 1* (GT_1) where we probed the notion of category structure induced by discrimination training by testing new stimuli spanning the entire stimuli space. As expected, the performance on this stage was lower than in the previous one (69.8% +- 2.8% of correct choices). In the second training stage, the *categorization training*, whenever mice reached 70% fraction of correct choices for two consecutive days, we added an extra pair of stimuli to the set, until reaching four per category (**Fig. 1C**). In this stage, mice spent 8.7 +- 1.2 training days reaching by the end of the stage a fraction of correct choices of 76.1% +- 1.8% with a discrimination index d’ of 1.4 +- 0.1. Lastly, mice underwent a second and *final generalization test* (generalization test 2, GT_2) with the same set of stimuli as in the first one, showing a performance of 70.2% +-1.53%.

To validate that mice were indeed using the stimuli frequency to perform the task, we fitted a logistic regression to predict choices on a trial-to-trial basis using stimuli tone frequency among other regressors (see Methods). The coefficient weighing the stimulus regressor (tone frequency) had the largest effect size and exhibited a significant upward trend throughout the training for the discrimination training stage (**Fig. 1E**, left, p<0.001, **Fig. S1C**) that then remained stably high during categorization training (Fig. 1E right, p=0.577, Fig. S1C). Consistently, there was a strong correlation between this stimuli coefficient and the corresponding fraction of correct responses (**Fig. 1F**, rho = 0.95 p<0.001, **Fig. S1D**). Together, these results validate the behavioral paradigm and confirm that mice relied on tone frequency to guide their choices, providing a robust framework to probe how hippocampal representations adapt to increasing category structure complexity and support generalization.

### Mice struggle to generalize for novel stimuli after discrimination but improve after explicit category training

We introduced two testing stages interleaved with the training ones to probe understanding of category structure. In these generalization tests we presented a set of stimuli spanning the entire stimuli space. We observed a drop in the total fraction of correct choices at the group level (**Fig. 1D**). To further study choice behavior at the subject level, we fitted psychometric sigmoid curves for each subject individually (**Fig. 2A, S2**) as described elsewhere [14] (see Methods), to characterize the decision function using tone frequency as the evidence variable.

**Figure 2:**
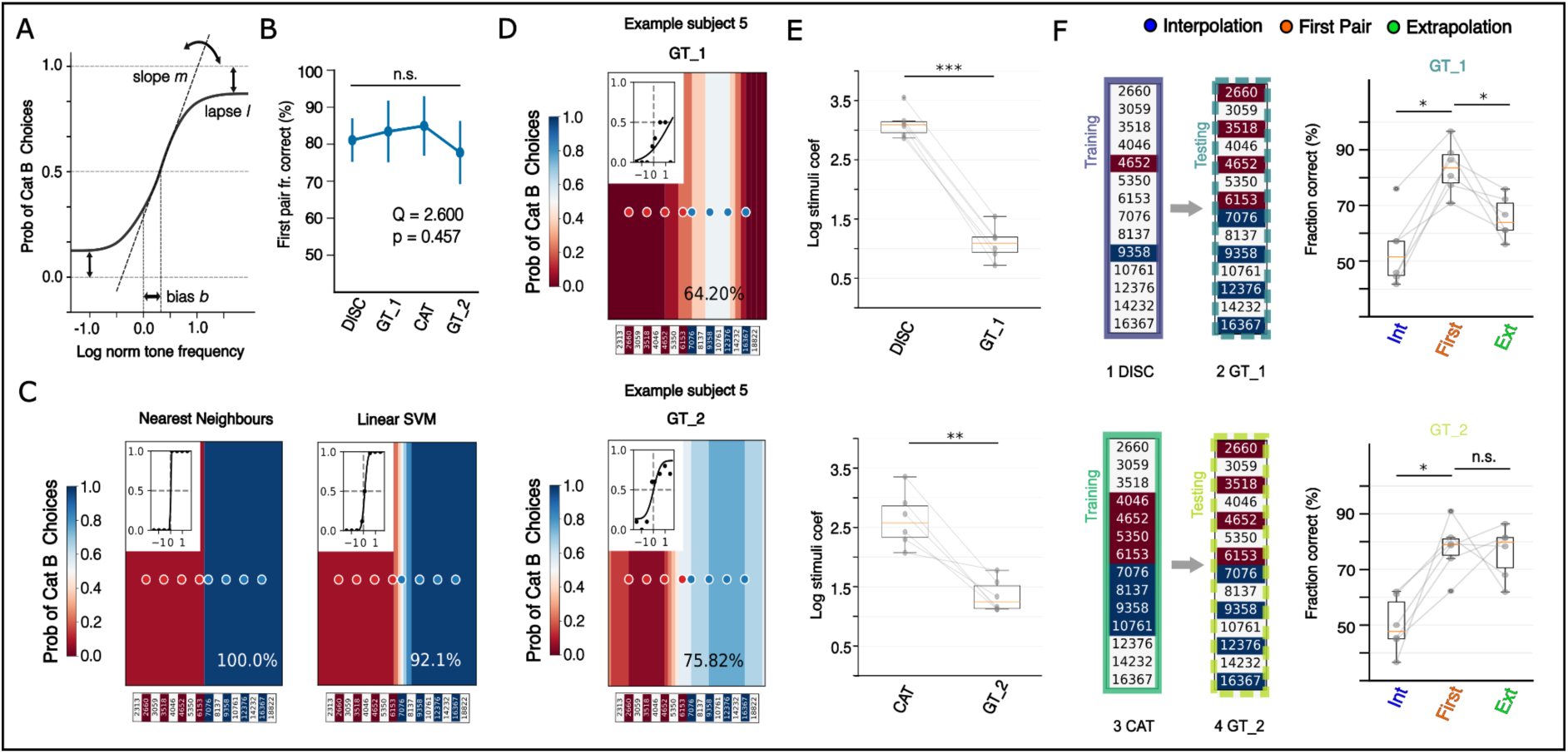
Mice behavior and performance in generalization tests. (A) Psychometric fit function consisting of the Error function with three parameters: slope, bias and lapse rate. (B) Fraction of correct choices for first pair of stimuli throughout stages showing stability (n=6, p=0.457, Friedman rank sum test). (C) Example decision plots (heatmap) and psychometric fits (inset) for two artificial classifiers (Nearest neighbors and Support vector machine) trained on discrimination stage and tested on generalization test 1, showing good fit and saturation of performance increasing with distance to boundary (See supplementary figure S2 for other classifiers). (D) Decision plot (heatmap) and psychometric fit (inset) for choices of example subject 5 in generalization test 1(top) and in generalization test 2 (bottom). Psychometric fit fails mostly at the edges of the stimuli space, most prominently at the high frequency end (See supplementary figure S2 for rest of subjects). (E) Logistic regression coefficient weighing for stimuli regressor drops in generalization tests compared to (top) previous discrimination stage (n=6, p=0.031, two-sided Wilcoxon signed rank test) and to (bottom) categorization stage (n=6, p=0.031, two-sided Wilcoxon signed rank test).(F) Fraction of correct comparisons (right) with corresponding stage schematics (left). Pair comparisons for fraction of correct responses between initial pair of stimuli (presented in discrimination stage) and inter- or extrapolation stimuli for (top) generalization test 1 (n=6, p=0.016,p=0.016, one-sided Wilcoxon signed rank test) and (bottom) generalization test 2 (n=6, p=0.016, p=0.5, one-sided Wilcoxon signed rank test).

This analysis uncovered a highly variable behavior among subjects that underperformed on generalization tests, despite being proficient on the corresponding previous training stages. However, the performance drop in the total fraction of correct choices was not explained by worse performance on the first stimuli pair (presented in the discrimination training), which showed no significant changes throughout the stages (p=0.457, Anova Repeated Measures) (**Fig. 2B**).

On the contrary, the most prominent struggle happened for novel stimuli, specifically those far from the boundary (extrapolation stimuli), particularly in the highest frequency end. This contrasts with the behavior assumed in psychometric fits and artificial classifier algorithms (trained in the discrimination pair and tested on the same test stimuli) (**Fig. 2C**), that expects that for stimuli far from the boundary the classification is easier. Indeed, the logistic regression model used to explain choices exhibited a significant drop in the stimulus coefficient (i.e. tone frequency) in both of the generalization tests, consistent with the behavioral struggle and decision-function maps (**Fig. 2E**). Specifically, for GT_1, failure to correctly classify the stimuli was more prominent for interpolation stimuli, i.e. stimuli within the range of the first stimuli pair and close to the boundary. However, as discussed above, extrapolation stimuli, i.e. stimuli at the edges of the stimuli space range, were not the easiest to classify, failing to reach an equivalent fraction of correct choices as for the first stimuli pair (p = 0.016) (**Fig. 2F**, top). Interestingly, in GT_2, after category training, performance on novel extrapolation stimuli improved (p = 0.5) (**Fig. 2F**, bottom), but performance on interpolated stimuli remained low, consistent with the profiles of psychometric fits that emphasize that stimuli close to the boundary are hardest to classify.

The stark difference between behavioral performance on extrapolation stimuli on GT1 (after discrimination training) and GT2 (after categorization training) suggests a change in behavioral strategy. The profile in GT1 is consistent with feature-based learning, that is, tied to the specific experienced stimulus value features (first pair of tone frequencies). Improved performance on GT2, by contrast, is compatible with a switch to rule-based learning (low vs high frequencies) (**Fig. 2F, S2**).

To explore the neural correlates of these behavioral changes we then turned to analyze corresponding neural activity recordings throughout the stages taking the last sessions for each as representative (see Methods).

### CA1 neurons exhibit a wide spectrum of heterogeneous single unit responses to trial events and variables

To image neuronal activities during task execution we injected all mice with a viral vector in the CA1 area to express a genetically encoded calcium indicator GCaMP8m (**Fig. 3A**) followed by a cannula implantation to enable chronic imaging of CA1 neural activity. After recovery, we then started habituating mice to the setup and training them on the most basic version of the task in a home-cage setting. We then used two-photon calcium imaging on selected days to record in head-fixed behaving mice (n=6) (Supplementary Movie 2). This allowed us to image the activity of neurons at a cellular resolution (n= total neurons; 306 +-12 CA1, (mean ± s.e.m.) simultaneously recorded per session) (**Fig. 3B, 3C**). To proceed with the neural data analysis, we made use of different events throughout the trial as time alignments. In this way, jittering of trial intervals leading to trial variability was taken into account (**Fig. 3D**).

**Figure 3:**
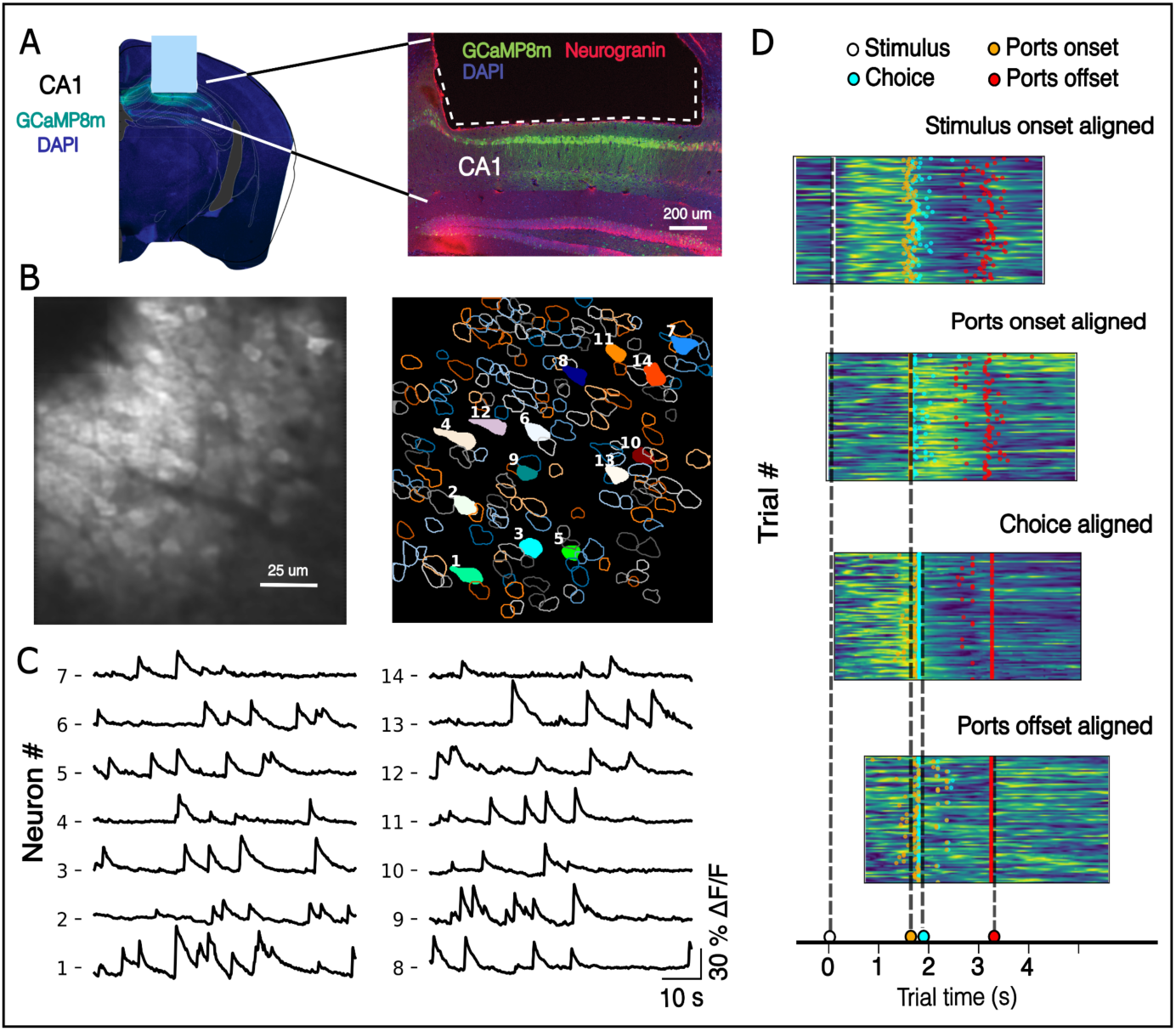
Calcium imaging and neural recording. (A) (left) Histology section showing grin lens implant and viral injection site. (Left) Calcium imaging was performed with a 2-photon microscope through a gradient index (GRIN) lens (light blue rectangle) that was implanted above the hippocampus CA1. GCaMP8m was genetically expressed in excitatory neurons through viral vector ssAAV injected in CA1 (see Methods). (Right) Close-up histology section showing expression of CA1 excitatory neurons with Green-GCaMP8m in the injection site under the grin lens (white dashed line), Red- Neurogranin, Blue-DAPI. (B) Example recording field of view of one frame (left) with corresponding segmentation of neuron cells extracted with CaImAn pipeline (right). (C) Corresponding normalized Ca +2 traces for numbered example cells in B. (D) Activity rasters across trials for 4 different example cells aligned to different trial events to account for jittering of intervals across trials.

To characterize neural activity, we first analyzed CA1 single unit responses by building peri-event time histograms (PETHs) aligned to stimulus and choice (licking) onsets. We then performed the so-called Zenith Event-based Time-locked anomalies test (ZETA test, [15]). This is a model-free test that identifies neurons with time-locked responses in relation to an event of interest (**Fig. 4A**). Neurons could show stereotypical responses in relation to stimulus onset (stimulus-modulated), to choice onset (choice-modulated), to both event onsets, or show no modulation at all (**Fig. 4B**). As previously reported [12,16], we found neurons showing significant activity tiling the events throughout the trial, in relation to the stimulus, choice and outcome onsets (**Fig. 4C**). However, we also found neurons that were responding selectively for certain event values, like for non-rewarded trials or even for specific trial types, showing interaction term responses (**Fig. 4D**).

**Figure 4:**
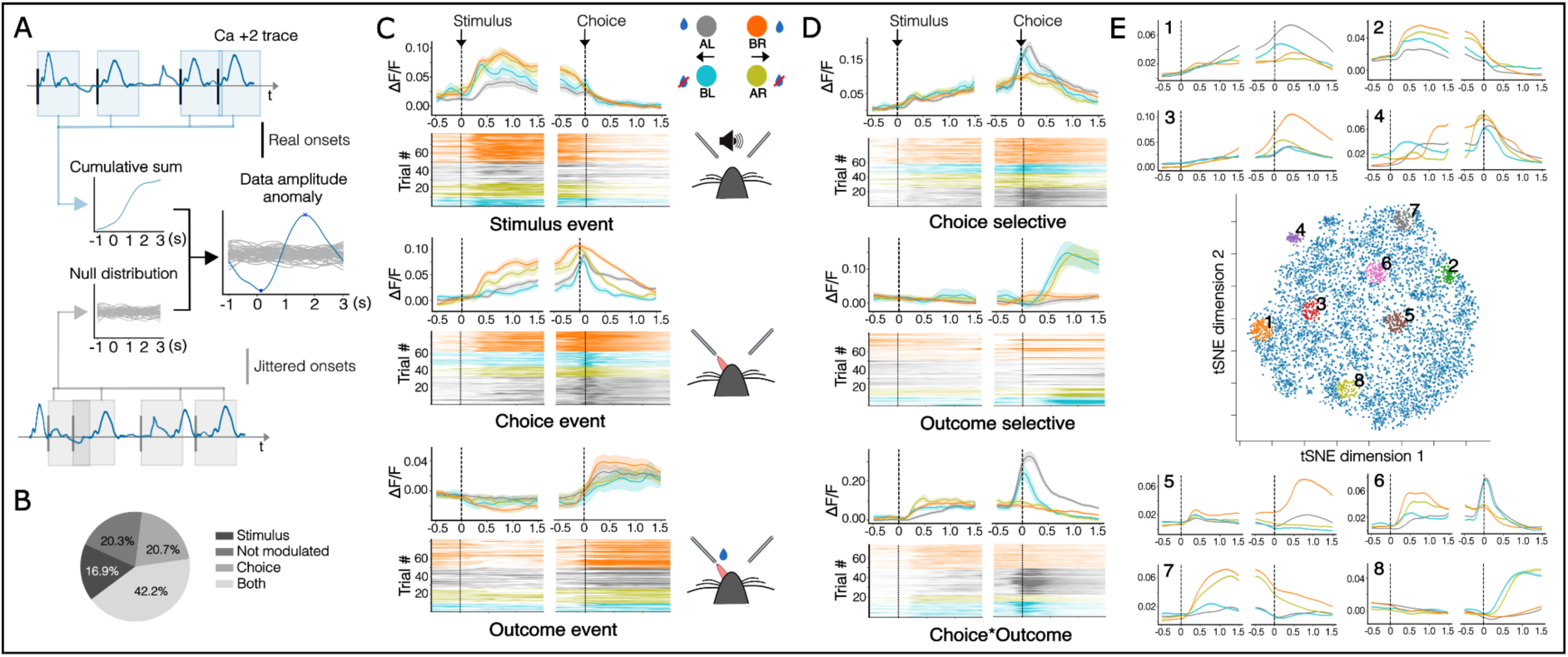
Overview of single neuron response profiles. (A) ZETA test method schematic. Single-trial responses following event onsets are tested against responses following randomly jittered onsets. (B) Proportion of modulated neurons according to the ZETA test on stimulus and choice onset alignment windows. (C) PETH and raster plot of example neurons responding to events in the trial with corresponding behaviour schematic. Tone stimulus presentation (top), choice licking (middle) and outcome (bottom). Trial types: AL: category A stimulus, left choice; BR: category B stimulus, right choice; BL: category B stimulus, left choice; AR: category A stimulus, right choice; (D) Example neurons with selective or mixed profiles. Choice selective (left choices) (top), outcome selective (no reward) (middle) and, choice (left) and outcome interacting (bottom). (E) t-SNE analysis on response profiles. Numbered PETHs correspond to average activity of the neurons selected in the corresponding colored neighbourhoods of the t-SNE map. 1: choice*outcome, 2: stimulus, 3: outcome*choice, 4: choice, 5: outcome selective, 6: choice selective, 7: choice selective, 8: outcome selective.

To address these response profiles, we performed a t-Distributed Stochastic Neighbour Embedding analysis (t-SNE) on condition-averaged responses. Specifically, for each neuron, we concatenated the average response to four trial types across two task events (stimulus and choice onsets) (see Methods, **Fig. S3B**). We observed a myriad of profiles resembling previously identified profiles (**Fig. 4E**), with no obvious clustering pattern (p>0.05, silhouette score, **Fig. S3D**).

In summary, CA1 neurons displayed highly heterogeneous and mixed selectivity profiles, consistent with a continuum of response types, aligning with results from previous work [16], rather than distinct functional subgroups. Given this diversity at the single-cell level, we next asked how the variables describing the different trial types were represented at the population level.

### Variables describing trial type are decodable from CA1 population activity

We therefore ran a series of support vector machine (SVM) linear decoders throughout trial time (**Fig. 5A**). We trained cross-validated, time-independent, SVM decoders on temporally aligned trials to two events: stimulus onset and choice licking onset (**Fig. 5B**). To create unbiased train and test sets, we sorted trials based on the variables describing each trial (category - A vs B, choice - left vs right and outcome - reward vs no reward), resulting in a total of four trial types. For each dichotomy, given the decoding variable, the other two were fully balanced through resampling, discarding sessions with not enough trials per type (see Methods). To increase statistical power, we used the pseudopopulation method to aggregate data across subjects as done similarly in other studies [13,17,18] (**Fig. 5C**).

**Figure 5:**
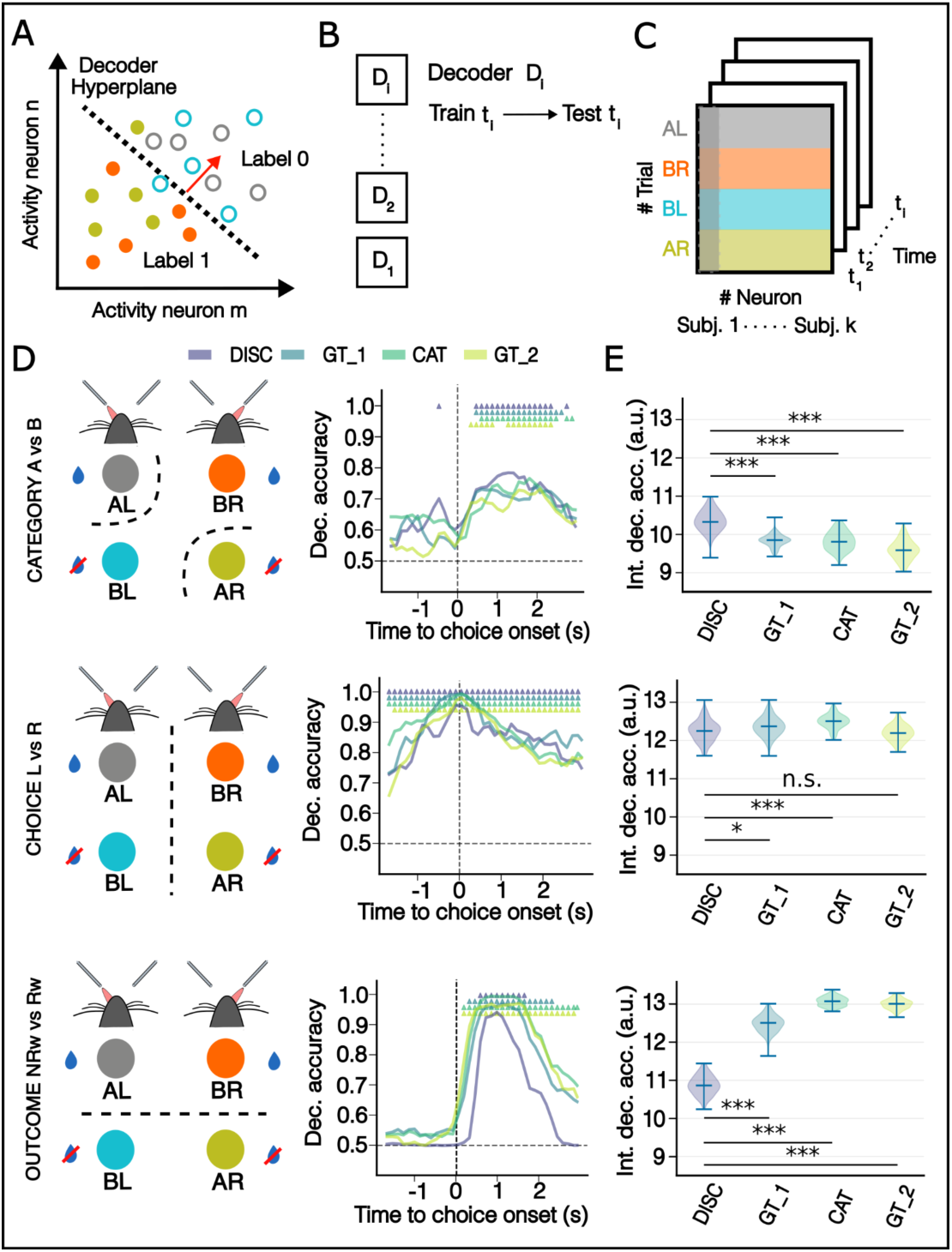
Decoding of main variables in task (category, choice and outcome) through trial time. (A) Schematic of SVM decoder separation of datapoints (trials) in neural space that creates a decoder hyperplane (dashed line) with corresponding decoder vector (red arrow). This example shows separation by classifying upon choice. (B) At each given timepoint in the trial we train a separate decoder and test it on the same timepoint. Each accuracy estimate is the mean of 100 cross validation runs and tested against a null distribution of 500 runs with permuted labels. (C) Decoder matrices are built by the pseudo population method to aggregate data across sessions and subjects, keeping critical trial conditions balanced. (D) For each row, decoding schematic (left) and decoding accuracy traces in trial time (right) for each variable. Trace colors correspond to the different stages in the experiment. DISC: discrimination, GT_1: generalization test 1, CAT: categorization, GT_2: generalization test 2. Colored triangles above traces label timepoints in which decoding accuracy departs significantly (one-sided, p< 0.05) as result from a percentile test upon the null distribution of permuted label runs and corrected for multiple comparisons taking the maximum of the percentiles across timepoints (see Methods). (E) Violinplots for integrated accuracy traces over trial time (bars indicate minimum, mean and maximum value of integrated decoding accuracy of each of the distributions of cross-validation runs) across stages. Overall trends are tested with Kruskall-Wallis test for category (top) (p < 0.001), choice (middle) (p < 0.001) and outcome (bottom) (p < 0.001) decoding. Stage pair comparisons (plotted with stars) are tested with Mann-Whitney test *: p < 0.05, **: p < 0.01, ***: p < 0.001.

In correspondence with the results at the single neuron level, we found that outcome and choice (licking side) were decodable in the CA1 population with high accuracy (>0.8, p<0.05, percentile test on permuted labels, **Fig. 5D**). Interestingly, the category variable, despite having no clear single neuron correlates, was also decodable above chance in all stages of training (>0.65, p < 0.05, percentile test on permuted labels **Fig. 5D**). Remarkably, neural activity decoding showed significant predictive accuracy for choice before the recorded behavioral event (**Fig. 5D** middle panel), extending as post-choice activity, overperforming decoders trained only on behavioral video footage (Mann Whitney U test, p<0.001, **Fig. S4A**). In contrast, the outcome variable was only decodable after the event, opposite to findings in other paradigms [12]. Critically, the category variable was also mostly decodable after choice (**Fig. 5D**), except in the discrimination and categorization stages where it was significant before choice for a few timepoints.

To understand the decoding accuracy changes across stages, we computed decoding accuracy (integrated over entire trial time). This analysis showed a small decrease for the category variable throughout stages, approximate stability for the choice variable and a prominent increase for outcome (p < 0.001, Kruskall Wallis test; Mann Whitney U test for stage pair comparisons with Bonferroni correction, **Fig. 5E**). Additionally, we computed the significance of the difference between them for each timepoint in trial time, across all possible stage pair comparisons (bootstrap percentile test, **Fig. S4B**, see Methods). The decoding accuracy for outcome in the post-choice window exhibited the most prominent effect size with a marked progression, consistent with the observation in the integrated decoding analysis (**Fig. 5E**)

Overall, these findings demonstrated prominent encoding of choice and outcome at the population level with a trial-time dynamics characterized by an overlap of significant decoding accuracies in the post-choice period. On the other hand, category information, despite being decodable above chance (also mostly in the post-choice period), was less prominent (maximum ∼0.75). Interestingly, the outcome variable exhibited the most marked stage effect mostly at the expense of increasing the time window in which the decoding accuracy was significant. To further understand the dynamics of representations and changes across stages, we performed a representation geometry analysis to unravel the robustness of these decodable variables.

### Factorized representations at population level for choice and outcome, not for category, emerge with category learning

Decoding accuracies observed in linear decoding analysis (**Fig. 5**), could be the result of different geometries in neural state space. Category decodability (A vs B) was less prominent and had apparently no direct single cell correlates, making it consistent as a result of high shattering dimensionality (ability to create a linear decoder in high dimensionality space for any given parsing of conditions, as defined in Bernardi et al [13]). On the contrary, choice (left vs right) and outcome (reward vs no reward) exhibited high decoding accuracies. Therefore, we hypothesized that choice and outcome would be represented robustly in “abstract format” [13], whereas category decodability would be a byproduct of the neural state space arrangement. To address this question, we examined the geometry of the representations that allowed for decoding of variables as seen in **Fig. 4**, using the cross-condition generalization performance (CCGP) geometry analysis as described in several studies [13,18,19].

We focused on the three main variables of each trial type: category, choice, and outcome. The four resulting trial types can be arranged in neural state space in several ways, depending on the dimensionality represented (considering the tuple of variables: category, choice and outcome, up to three) and the relative positioning of states, to support successful linear decoding (**Fig. 6A**). We characterized these arrangements by analyzing the cross-condition generalization performance (CCGP) of each variable, splitting it for each of the other two (decoding accuracy on x with split training and testing according to y: Cc_x_y_; see Methods). This analysis allows to test whether the hippocampus encodes a variable in a robust format, enabling generalization across conditions given by the rest of variables. Although the task structure ensures by design that any two variables (from category, choice and outcome) uniquely determine the third, mice may not necessarily exploit this dependency, nor is it clear which variables would dominate the hippocampal code, especially considering that all three are linearly decodable from population activity. For this reason, we examined the representational geometry of all three variables as a whole for completion.

**Figure 6:**
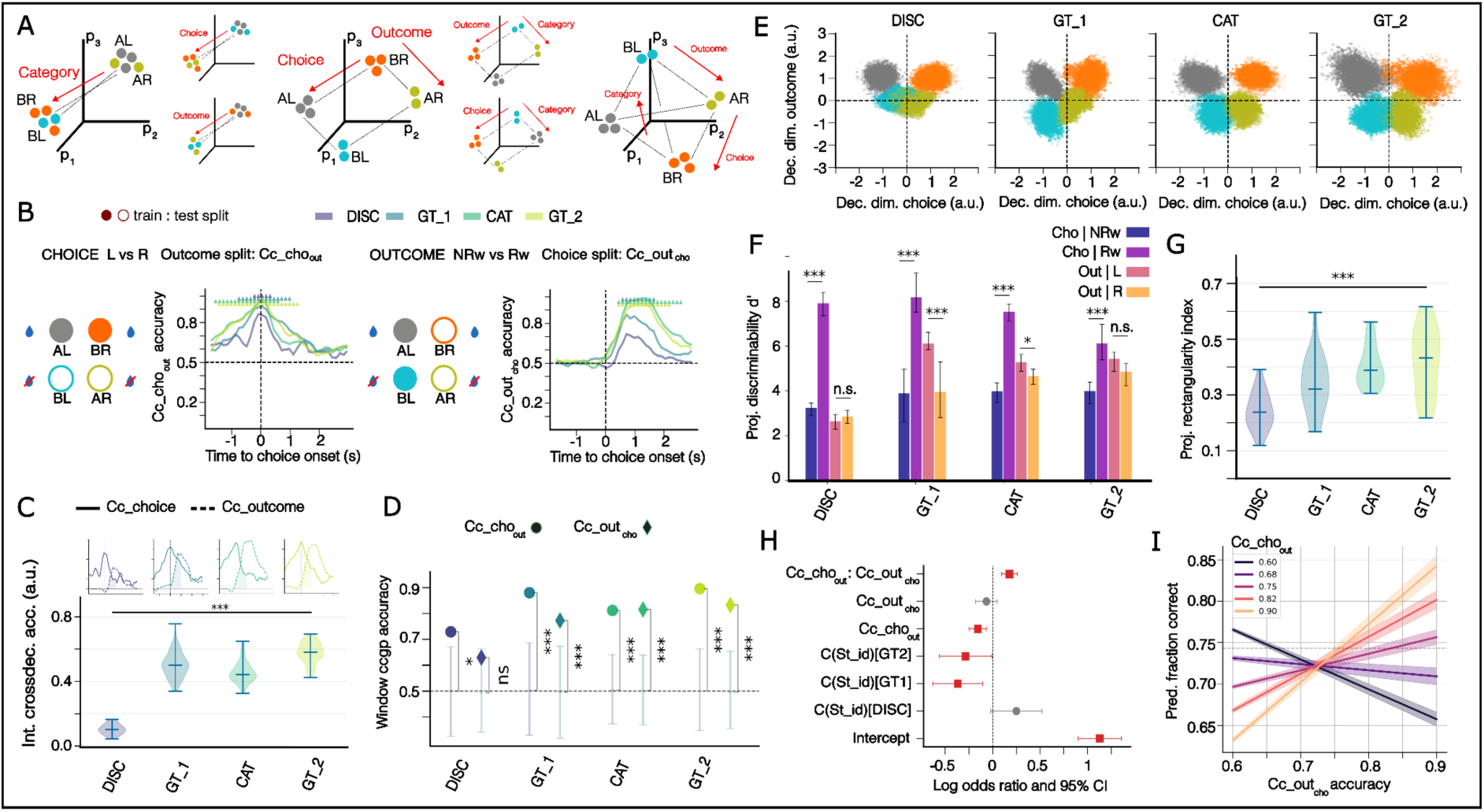
Geometry decoding analysis of main task variables. (A) Schematic showing hypothesis of representations geometry in neural state space taking the three main variables: category, choice and outcome. Possible geometries differ in dimensionality and in the specific arrangement of the different trial type conditions. Red arrows and labels indicate variables corresponding to the decoding axis. (Left) One-dimensional geometries where only one of three variables is robustly decodable. (Middle) Two-dimensional geometries with two orthogonal decodable variables. (Right) three-dimensional arrangement (only one possibility) would allow any successful decoding of all three variables. (B) Cross-condition decoding schematic and corresponding CCGP accuracy trace throughout trial time for each of the stages for choice with outcome split (Cc_cho_out_) (left) and outcome with choice split (Cc_out_cho_) (right). See Fig. S5 to see the rest of cross decoding splits. Significance is computed against null distribution built with 500 permuted label runs and corrected for multiple comparisons taking the maximum of the percentiles across timepoints. (See methods). (C) Integrated cross-decoding accuracy overlap of Cc_cho_out_ and Cc_out_cho_ integrated for post-choice window (shaded area) at different stages. Violin plots show distribution of cross-validation runs (bars correspond to minimum, mean and maximum accuracy values on the distribution) (Kruskall-Wallis test, p < 0.001). (D) Cc_cho_out_ and Cc_out_cho_ computed at the post-choice window across stages. Discrmination is the only stage where both are not significant from chance (p-values computed against null distribution of permuted label runs) *: p < 0.05, **: p < 0.01, ***: p < 0.001. (E) Neural state space projection onto the plane given by the decoding vector for choice and the decoding vector for outcome. Each datapoint corresponds to the population activity corresponding to one trial in the post-choice window, color-coded according to trial type. (F) Discriminability of choice or outcome computed on the projected neural state space across stages, conditioned on specific values of choice (left or right) or outcome (rewarded or non-rewarded) (paired t-test on cross-validation runs). (G) Rectangularity index (see Methods) computed on the projected neural state space in E, exhibiting a marked increase across stages (One-way Anova test, p < 0.001). (H) Forest plot of predictor coefficient values in a log scale for the selected binomial regression model with corresponding 95% confident interval. Red predictors are significant. (I) Interaction term effect plot showing increase of slope fraction of correct choices vs Cc_out_cho_ with increasing values of Cc_cho_out_.

We found that cross-condition decoding splits (Cc_cat_cho_, Cc_cat_out_) for the category variable led to accuracies significantly below chance level throughout the trial (anti-generalization) (double-sided percentile test on null distribution of permuted labels, **Fig. S5A**), revealing that the decoder was actually capturing information from either choice or outcome from the training trials. For example, if a decoder trained to distinguish category A vs. B with outcome split (using AR vs BL, non-rewarded trials) actually captures choice information (R vs. L) instead, it will predict the opposite category label when tested (AL vs BR, rewarded trials). This cross-condition decoding analysis aligned with our hypothesis that category information was not represented robustly but instead was decodable as a byproduct of the representation geometry. In contrast, choice and outcome exhibited high CCGP values for corresponding cross-splits against one another (Cc_cho_out_, Cc_out_cho_), showing robust decoding (**Fig. 6B**).

Simultaneous significant accuracy values for Cc_cho_out_ and Cc_out_cho_, as observed during the post-choice period, would enable disentangled readout of choice and outcome by other areas, independently of the exact combination of variable values (left or right choice, rewarded or not) present in the trials. These representation geometry findings ruled out 1-dimensional arrangements explained in **Fig. 6A**, where only one variable would present with high cross-condition decoding. To explore whether this disentangled format of choice and outcome was relevant in magnitude or even at all related to the complexity of the task, we tested for stage effect by computing an overlap of decoding accuracies of Cc_cho_out_ and Cc_out_cho_ (integrated in trial time over a post-choice window). Stage differences were most striking between the discrimination stage and the rest of stages showing a significant overall progression (Kruskall-Wallis test, p < 0.001, **Fig. 6C**). Furthermore, cross-decoding accuracies (Cc_cho_out_, Cc_out_cho_) computed solely as post-choice window averages (instead of integrated) were simultaneously and significantly high, except in the discrimination stage where Cc_out_cho_ accuracy failed to reach significance (percentile test on null distribution of permuted labels, **Fig. 6D**) aligning with our previous result. Interestingly, discrimination training is the only stage where the complexity of the underlying category structure has not been presented yet to the mouse.

Moreover, neural population activity in the post-choice window projected onto the decoding axis of choice and outcome (**Fig. 6E**) showed that representation geometry is increasingly factorized (more rectangular in shape) with improvement in classification discriminability (d’ prime) (**Fig. 6F, 6G**). Again, this property is not present in the discrimination stage, despite having a significant linear decoding (for both choice and outcome, **Fig. 5D**) and proficient behavioral performance, reflecting on the nature of the category complexity presented. Instead, after discrimination, the results point towards a representation geometry that increasingly factorizes choice and outcome, as confirmed in the projected data (**Fig. 6E**), implying representation geometries with dimensionality greater than two (in regard to the three main variables). This is further supported by state space simulation results, that when constrained to observed decoding properties of data, (**Fig. S5C, S5D**) rule out 1-dimensional arrangements, matching our findings (see Methods).

In summary, our geometry analysis revealed that CA1 population codes for choice and outcome became increasingly factorized across stages during the post-choice window, forming an abstract, disentangled format that was absent during simple discrimination but emerged when exposing the mice to a higher complexity on the category structure. During discrimination stage, cross-decoding accuracy results showed only choice was robustly encoded, fitting onto a 1-dimensional geometry governed by a choice axis. However, as soon as more stimuli per category were presented and the combination of choice and outcome did not uniquely determine the tone, we observed a progressive increase in outcome discriminability independent from choice (**Fig. 6E, 6F, 6G**), adapting the encoding to a factorized choice-outcome representation (at least 2-dimensional).

We then hypothesized that as the task gets more complex, this disentangled readout would be advantageous to solve the task. To test whether indeed this factorized format provided a computational advantage for behavior, we next examined how the geometry of choice–outcome representations related to task performance.

### Disentangled representation geometry of choice and outcome explains behavioral performance

Finally, to examine whether this “abstract” encoding format of choice and outcome was related to behavioral performance in the task, we fitted a binomial regression model to predict the fraction of correct choices using Cc_cho_out_ and Cc_out_cho_ as regressors computed for each subject at each stage (see Methods). To avoid biasing the results on which regressors turned out as significant in the most parsimonious manner, we ran several models with and without the interaction term of the two regressors (Cc_cho_out_:Cc_out_cho_) and extra categorical regressors accounting for data structure (see Methods). Next, we selected the best model by minimizing the Bayesian Information Criterion (BIC). The selected model: (Nr correct, Nr incorrect) ∼ C(stage id) + Cc_cho_out_ + Cc_out_cho_ + Cc_cho_out_ : Cc_out_cho_) results with a BIC of 149.24 and deviance explained of 58.90 and shows robustness of results with no qualitative changes when running the model without points with high leverage or outliers (**Fig. S6E, S6F**). Regressor for discrimination stage (first level of stage regressor, i.e. stage id) weighted positively (although not significant), whereas regressors for generalization tests had a significant negative coefficient. This aligns with the observed behavioral effect of discrimination being the stage with best performance and generalization tests constituting a struggle.

Remarkably, the resulting model included the interaction term (Cc_cho_out_: Cc_out_cho_) and exhibited a positive and significant coefficient in explaining total performance (0.1754 p<0.001, **Fig. 6H**), significantly enhancing the reduced model without interaction term (p<0.001, Likelihood ratio test, LRT, **Fig. S6G**). Specifically, the interaction term would start to have a net positive effect on the fraction of correct choices in the higher range of the decoding accuracy (>∼0.7) for both Cc_cho_out_ and Cc_out_cho_ (**Fig. 6I, S6A**), that corresponds to a factorized (choice-outcome) geometry. As demonstrated in simulations, constraining to high values of cross-decoding accuracy precisely rules out 1-dimensional arrangements (**Fig. S5D**). On the contrary, any of these cross-decoding accuracies, taken independently, contributed negatively to performance (-0.1556 std.u. for Cc_cho_out_, p=0.001 and -0.0660 std.u. for Cc_out_cho_ p=0.256) (**Fig. 6H**). This validates our hypothesis, since these results imply that non-factorized geometry (absence of high cross-decoding accuracies for both choice and outcome), as opposed to a factorized one, results in a worse performance.

Together, our results show that the emergence of a factorized geometry in CA1 (of choice and outcome) that allows for disentangled readout explains improved behavior performance, as opposed to representation arrangements where choice and outcome are linearly decodable but do not generalize across trial types (non-factorized geometry).

## Discussion

We characterized hippocampal activity of a population of CA1 neurons as mice trained on a novel categorization paradigm. Specifically, we introduced a novel training paradigm consisting of different training stages with interleaved generalization tests to probe category understanding. This allowed us to explicitly test the generalization bias induced after discrimination training and compare it with the one after categorization training. Furthermore, these generalization tests included extrapolation stimuli (beyond the range of the stimuli presented in discrimination), in contrast to previous work that only included further training with interpolation stimuli [20,21]. This enabled us to check if generalization performance was actually tied to experienced tones (feature-based) or, on the contrary, mice were able to classify novel tones according to the rule (low vs high frequencies). We found that initially, after discrimination training, most mice struggled to classify these novel extrapolation stimuli. They exhibited no saturation of decision at the edges of the stimuli space as opposed to sigmoidal fits used in psychometric analysis [14,21]. In other words, despite novel stimuli being farther from the boundary in comparison to the trained stimuli, mice did not classify them better. This result aligns better with feature-based rather than rule-based generalization, as discussed in previous work showing that rodents and other animals tend to generalize upon experienced features and, humans, by contrast, tend to generalize upon rules [22]. However, after explicit category training in the second generalization test, performance improved on the novel stimuli that were far from the boundary. This suggests that a switch to rule-based generalization may have occurred or, alternatively, that feature-based representation of the categories (if considered as a likelihood distribution of category membership upon tone frequency) expanded after categorization training, *covering* novel extrapolation stimuli more successfully (**Fig. S2**).

By staging discrimination before categorization, we isolated category-structure demands from trial-structure learning, unraveling critical changes in the population code regarding representation geometry. During discrimination, mice were not exposed to the full category structure while they learned the dependencies and structure of trial events. On the other hand, in categorization training, trial structure was no longer novel and difficulty lied purely on learning the membership of more tone examples. In other words, category became a latent variable to be inferred. From neural population activity we could decode variables like category, choice and outcome (critical in describing the trial type), specially in the post-choice period and regardless of the stage. However, we determined relevant stage differences in decoding accuracy, in particular, the time length of the window in which decoding was above chance. Specifically, the outcome variable showed the most striking progression across stages (**Fig. 5D, 5E, S4B**). Moreover, our representation geometry analysis revealed that discrimination stage exhibited a lack of factorized choice-outcome representation (high Cc_cho_out_ but non-significant Cc_out_cho_) in comparison to categorization training where both of these cross-decoding conditions (Cc_cho_out_ and Cc_out_cho_) were simultaneously significant in the post-choice period, corresponding to a disentangled representation (abstract, factorized format for variables [13]). This change in representation geometry emerged at the expense of increasing outcome decodability along with a striking rise in choice decodability among non-rewarded trials (**Fig. 6E, 6F**) that became more informative across stages. Importantly, differences of trial encoding regarding choice and outcome between discrimination and categorization stages appeared even though trial structure and event dependencies were unchanged and began, reassuringly, at the first generalization test (**Fig. 6G**), suggesting that the exposure to the complexity of the underlying category structure and not choice policy or performance (that were alike) (**Fig. 1D, 1E**), influenced the representation change. During the first stage (discrimination), category assignment is a one to one mapping with tone frequency and vice versa. In this setting, we concluded that being sensitive to salient trials (only rewarded) with high choice discriminability seems to be enough for mice to learn the category assignment. However, as soon as the complexity of the category structure was displayed (from generalization test 1 onwards), this unique mapping breaks and category membership becomes a latent variable. Hence, we argue that having access to trial episodes in a choice-outcome disentangled format to retrieve trial information, regardless of exact experienced choice-outcome combinations, becomes a computational advantage.

By running a binomial regression model, we explicitly demonstrated that this change of representation to a factorized geometry (choice-outcome) indeed relates to an advantage in solving the task. We showed that this disentangled format (represented by a high interaction term of cross-condition decoding) positively contributes to behavioral performance. In fact, our results give further experimental support to other work in describing this representation format as critical to enable generalization abilities. Disentangled format of variables has been found to describe representation of task-proficient animals and humans [13,19]. Taken together these findings imply that episodic encoding of trial events would be critically affected by the complexity of the category structure presented, despite preserving the same trial event structure throughout the paradigm. Hence, episodic trial encoding, as means to create a state map (cognitive map), seems not only to be determined by relevant events and their relationship in the trial structure, as described in previous work [16] but also, potentially by computational demands that might change with task complexity, e.g. category structure.

Interestingly, we found that categorical information itself was not robustly represented in the hippocampal population. The decoding accuracy for category was in general not significant after stimulus presentation (barely one significant timepoint, in discrimination stage, **Fig. 5D**), suggesting that this information was not prominent in CA1 population activity after stimulus presentation (evoked response). This was the case even for proficient mice in discrimination stage (most proficient stage) where category decoding is equivalent to stimulus identity decoding. Moreover, the geometry analysis revealed that even the category decodability (moderate, maximum ∼0.75) in the post-choice window was not robust. Neural state simulations constrained by our experimental decoding properties showed that such above-chance accuracy can arise without a dedicated axis of the neural representation, effectively missclassifying one of four trial types (∼75 % correct). We therefore interpret this category decodability as a byproduct of geometrical arrangement of choice and outcome. These results support the idea that without informative transition structure (on category labels or feature dimensions serving category assignment), CA1 does not treat category as a context variable like for example, in Mishchanchuk, et al [12], precluding the encoding of the latent variable of category, despite being proficient on it (especially in discrimination stage). The opposite is described for variables found to be encoded in CA1 hippocampus (if predictive in a sequence of events) like in spatial navigation tasks where reward is *anchored* as an event in certain spatial locations (with reliable transitions) as described in Sun, et al [16], and in non-spatial paradigms, like in Aronov et al [4] where again, stimulus dimension (tone frequency) is presented in a reliable sequence. In our paradigm, by contrast, there is no structure within or across trials for categorical information nor a structured event sequence that would serve as a scaffold to map it into.

Multiple exciting directions for future research emerge from our results. Although our findings give insight on how trial episode neural representations may change with complexity of the task and not only with event trial structure, there is still further work needed to analyze the causality of these representation geometry changes, as being top-down induction from other areas as suggested by human experiments [19] where representation geometry of critical variables factorizes (disentangles) with verbal instructions given to the participant or if they arise bottom-up from local processes, or even as a combination of these. Moreover, another line of research would involve theoretical modelling to explore the computational advantage of this factorized format when learning a category structure. This will help to understand which down-stream computations required for category learning are precluded or enabled by this computational format of trial episodic memories provided by the hippocampus.

In conclusion, by combining large-scale calcium imaging with a multi-staged non-sequential categorization task, we show that CA1 representations are not tied only to pure event dependencies but rather adapt flexibly to the computational demands of category learning. Our findings reveal that the hippocampus reshapes trial event encoding into a disentangled format that supports generalization, highlighting its critical role in bridging episodic experience with category structure (abstract) learning.

## Methods

All procedures were approved by the Veterinary office of the Kanton Zurich (Cantonal Veterinary Office Zurich).

### Subjects

Experiments were performed on male C57BL6/J mice (Charles River, Germany), between 7 to 18 weeks old during the behavioral experiment. Mice were given at least 1 week period to acclimatize to the new housing conditions prior to the first surgery. After surgical recovery (at least 1 week), animals were housed by groups of up to 5 littermates in individually ventilated cages (IVC, Allentown T1800), in an inverted 12 h light/12 h dark cycle room (lights on from 7:00 pm to 7:00 am) and were provided food and water ad libitum. Mice had access to a running wheel and other enrichment material such as a mousehouse, bedding material, crinkles and a tunnel. During the experiment, mice entered a water restriction schedule, (at least 1 week before the start of the experiment) with access during the behavioral day sessions to ∼ 300-800 uL depending on performance and an extra ad libitum access to water for 1 continuous hour per day.

### Surgical procedures

#### Anesthesia

For all surgical procedures, preemptive subcutaneous Buprenorphine (Bupaq; Streuli, 0.1 mg/Kg) and Carprofen (Rymadyl;Zoetis, 4mg/Kg) were administered 30 minutes prior to anesthesia. Induction consisted of either a intraperitoneal Ketamin-Xylazin cocktail (Ketanarcon; Streuli, 75-90 mg/Kg / Xylazin; Streuli, 5-8 mg/Kg), or gaseous Isoflurane 4% (Attane, Piramal). Mice were subsequently mounted onto a stereotactic frame (Kopf Instruments). and the eyes were covered with ophthalmic ointment (Hylo Night, PharmaMeda). Anesthesia maintenance was achieved with concentration of Isoflurane (1-2%) delivered with (0.6-0.8L/min) medical oxygen (Pangas UN 1072, Conoxia) through a face mask. Body temperature was kept at 37 degree Celsius with a heating pad and a temperature controller throughout the surgery.

#### Stereotactic viral injection

After removing hair from the scalp and sterilizing it with Betadine, the skull was exposed with a single incision along the midline after application of subcutaneous Lidocaine (4mg/kg). After removing the connective tissue with sterile cotton buds, a small hole was drilled in the skull at the coordinates (1.8 mm posterior, 1.6 mm lateral, 1.5 mm depth relative to bregma) using a stainless steel burr attached to a miniature drill. 500nl of an adeno-associated virus were injected intracranially, driving the expression of the calcium indicator GCaMP8m via the CamKII promoter (ssAAV-9/2-mCaMKII-alpha-jGCaMP8m-WPRE-bGHp(A), VVF, UZH virus fabricat, 7.8*10^12 copies, titer 1:4) with a micropump (UMP3UltraMicroPump, World Precision Instruments) at a speed of approximately 50 nl/min. After injection, the needle was left in place for 10 minutes before being slowly retracted to avoid back-spill of the virus and finally the skin was closed using a running suture (silk braided suture 3-0, B Braun). At the time of the viral injection surgery, mice were 8-10 weeks old.

#### Cannula and headbar implantation

Before surgery, a cannula implant was prepared by attaching a 3 mm diameter glass coverslip (CS-3R-0 Cat:64-0726, Warner Instruments) to the edge of a 1.5mm diameter - hollow sheet metal cylinder, using a UV-curing optical glue (Norland optical adhesive 81, Norland Products). Then the extra glass cover was grinded down with a grinder (Ultrapol, Ultra Tec) to match the cylindrical sheet diameter and left to fully cure at room temperature for a minimum of 24 h.

Once the proper anesthesia depth was confirmed and the animal was mounted into the stereotaxic frame, the skull was exposed, dried and scraped with a scalpel to improve adherence of the head plate. A 1.5 mm diameter craniotomy, centered at the previous injection site was performed using a dental drill. Subsequently, a durotomy was performed using micro-tweezers (11273-22, Fine Science Tools) and with gentle aspiration the cortex above CA1 was removed until the corpus callosum fibers were visible. After hemostasis, the cannula implant was introduced in place and secured with light curated glue (Scotchbond, 3M). A custom-designed titanium head plate was aligned and centered at the midline, and attached with a first layer of the same light curated glue (Scotchbond, 3M) and then secured with dental acrylic (Paladur, Kulzer). One anchoring screw was fixed to the skull in the contralateral side to ensure a proper adherence of the cement and the head bar.

Lastly, a RFID tag (125kHz, Sparkfun) was placed subcutaneously at the posterior part of the neck, ensuring that neck mobility was spared.

#### Post-surgery recovery and analgesic regime

Mice were allowed to recover after surgery for a minimum of 30 minutes in a wake up cage placed on a heating plate (37 degrees) before they were returned to their home cage. For 3 days after each surgical procedure animals received Buprenorphine s.c. (Bupaq; Streuli, 0.1 mg/Kg) every 6 h during the dark cycle and in the drinking water (Bupaq; Streuli, 0.01mg/mL) during the light cycle, as well as Carprofen s.c. (Rimadyl; Zoetis, 4mg/Kg) every 12 h.

#### Preparation of animals for behavioral experiments with recordings

After cannula and head-bar implantation a period of at least 4 weeks before testing viral expression levels was given. Approximately one week before imaging sessions we inserted the gradient index (GRIN) lens (CR3530-2, GT-IFRL-100-101027-50-NC; GRIN-Tech) into the guide tube fixing it to the cannula with optical glue (Norland optical adhesive 81, Norland Products)

### Validation of imaging methodology

#### Perfusion

After recordings were completed, mice underwent terminal anesthesia with Pentobarbital (Esconarkon; Streuli, 200 mg/Kg). We transcardially perfused all mice with PBS followed by cooled paraformaldehyde (4% PFA). Next, we fixed the entire head in PFA for an extra week.

#### Verification of cannula implant

We then removed the entire brain and used a Vibratome (VT2000s, Leica) to cut 50 um thick coronal brain slices and stored them in PBS and mounted all slices on microscope slides and acquired large field-of-view images with a standard fluorescence microscope (Z16, Leica) using two channels: FITC for GCaMP8m and UV for DAPI. Finally, we used a reference brain atlas [23] to check the position of the grin lens in the recordings for each subject.

### Behavioral procedures

#### Behavioral setup

We trained animals on an auditory binary categorization task. Following a minimum of 7 days of recovery after head-bar and cannula implantation surgery, mice were water-restricted to up to 85% of their ad-libitum weights. After at least a week of water-restriction and habituation to manual handling by the experimenter and familiarization with the home cage and training chamber, behavioral training for the binary auditory categorization task began.

All behavioral experiments were performed in a customized 3D-printed training chamber, attached to the side of an adapted standard home-cage (small, Mouse 500). The end of the training chamber was equipped with a latching system in which the mice learned to head-fixate themselves, aligned under the two-photon microscope. In front, two retractable stainless steel licking ports were located, mounted on stepper motors (1204, Pololu robotics). Licking spouts were connected to a closed-loop sensor circuit [24] for lick detection in combination with an aluminum floor in the training chamber. Auditory stimuli were presented to mice subjects via a speaker in front. Experimental events were controlled and recorded using custom Python scripts running on a Raspberry Pi.

All training and recording sessions were 35 minutes long consisting of approximately 100 trials depending on mice performance. For each trial after stimulus presentation, a delay period lasting up to 1.2s was delivered. After this delay, the licking ports moved into reach for the response window where the mouse could lick, lasting for 2 s. The first lick on the lick spout within the response window determined the response for the trial, resulting in immediate delivery of a water reward if correct or a timeout if incorrect. The trial finished with an inter-trial interval of 3.5-4.5 ± 0.15 s leading to the next trial. All intervals (delay period and inter-trial period) except for the stimulus presentation and the response window were randomly jittered to reduce time based prediction of events. Auditory stimuli space consisted of a discrete set of pure tone sounds ranging from 2000Hz to 20000Hz in frequency with a power-law spacing with a total of 16 tones, equally spaced.

#### Training stages

Prior to data collection, mice went through a pre-training period to engage voluntarily with the head-fixation system. After initial handling and getting familiarized with the home-cage and the training chamber, all mice were placed on the home-cage and left at their leisure to enter the training chamber and collect water ad libitum by licking the water spouts, at first without head-fixation. Next, head-fixation was introduced for increasing times allowing water consumption by operant licking from the spouts without any trial structure. Once mice appeared calm and confidently lick enough water during these fixation periods, the full task structure was introduced. The first stage of the paradigm consisted of discrimination. To promote engagement and avoid bias in licking, first sessions included 80% of passive trials that delivered a water drop from the corresponding water spout according to the stimulus presented. If mice showed less than 15% response rate difference between the ports, the fraction of passive trials diminished progressively until all trials were dependent upon mice response. Next, the grace period (interval between stimulus offset and response window onset) would be shaped from the initial 600ms up to 1.2 s depending on reaching at least 70% performance on two consecutive days. Imaging recording commenced in this stage after mice met the performance criterion with low bias and the final length for the grace period. After proficiency in this last step of the discrimination stage, mice were then introduced to the first generalization test. This consisted of just one session (to reduce likelihood of learning) to probe understanding of the category structure by presenting a larger set of stimuli per category spanning the stimuli space. Animals then entered the categorization training phase where a small subset of the stimuli space was presented until they reached the ‘expert’ level with the consistent performance of over 70% for at least two consecutive sessions. Lastly, mice were tested again for a second generalization final session equivalent to the first in the available stimuli set.

#### Calcium imaging during mouse learning behavior

Calcium imaging was performed with a two-photon microscope with a resonant scanner (Scientifica). The light source was a pulsed femtosecond laser tuned to 940 nm (Spectra-Physics) and the objective was a 20x (Olympus, 0.45 NA). Functional images detecting fluorescence signals of GCaMP8m were acquired using ScanImage (Vidrio) software. Imaging was done in one plane with a 2x magnification over a field of view of 500 mm x 500 mm (512x512 pixels), with a sampling rate of 30 Hz through the GRIN lens. (Supplementary video 2) The average laser power under the objective ranged from 50 to 80 mW. Note that the laser power was higher than for imaging through a conventional cranial window due to power loss over the grin lens, hence the power under the objective served as an upper bound on the power delivered. The same imaging plane was found on successive days by visual comparison to saved imaging, aiming for the same position as the stored absolute coordinates of microscope position from previous days.

### Extracting neural activity from calcium imaging data

#### Denoising

The first step in the preprocessing pipeline was denoising of the imaging stacks. Before performing the signal extraction on the imaging stack, each session was denoised using the DeepCad-RT library [25]. Training of the denoising models was done for each of the mouse subjects separately. 2000 frames were collected from the baseline recording (at the beginning of the session) of the first imaging session to train each subject model. Next, the rest of the sessions of each subject were processed with the resulting model.

#### Cell extraction for individual sessions

To automatically identify individual neurons in the calcium imaging movies of a given imaging session, we used a well-established cell extraction analysis pipeline (CaImAn, Python [26]). Imaging data was first processed to remove translation and rotation motion artifacts via the NoRMCorre algorithm [27]. Next, source extraction was performed using the constrained non-negative matrix factorization (CNMF) [28].

#### Session alignment

To be able to track cells across imaging sessions we implemented an alignment procedure with a combination of custom written Python code based on the phase-cross-correlation Skimage library function and the cell matching algorithm used in the register-multisession function from the CaImAn library. We first constructed cell templates for every session using the motion-corrected average image obtained from the cell extraction algorithm. Next, we estimated the shift among them by analyzing the highest peak in the cross-correlation landscape between any pair of images in the Fourier space, getting a common field of view (FOV) across sessions. This customized code was then fed to the multiregistration-rois function of the CaImAn library based on the Hungarian algorithm that solves the assignment problem identifying pairs of cells whose centroids match within a certain neighborhood. The resulting method matched about 15-75% of the originally identified cells of a single recording depending on the distance in time between recordings.

#### Post-processing and validation

Validation of automatically extracted neurons was completed with a semi-manual process to further determine the quality of each component. When running the CaImAn pipeline, a threshold for the SNR of the traces was specified to a value of 2 as a first criterion of quality. Thus, all components with SNR below 2 were labelled as rejected. The remaining components underwent additional screening from a support vector machine (SVM) algorithm to classify cells as accepted or rejected. To train this model, 5% of identified cells from all experimental imaging sessions were manually validated. Then, a series of hand-crafted features were computed for each cell component. These features consisted of various morphological like cell diameter, perimeter, area, compactness and trace signal characteristics: trace variance, first component of Fourier transform and asymmetry of trace. The resulting SVM classifier was evaluated on the ground truth dataset (manually validated) resulting in a false positive rate of 2% and a total acceptance rate of 76% of all identified cells accepted automatically by CaImAn.

### Quantification and data analysis

#### Behavior analysis

Performance was computed as the fraction of correct trials out of all active trials in which the subject had a response. These trials were the majority for trained mice even when responding incorrectly since the engagement levels were high and bias decreased progressively and was enforced to be low to progress through the stages of the experiment. On the other hand, discrimination (d’) was computed as a more precise parameter to demonstrate learning of the task. It was computed as described elsewhere [29] adapting it to two licking ports. Conditional probabilities in analyzing response policies were computed on fraction of condition trials over total of active trials, disregarding non responsive or passive trials.

#### Logistic regression analysis

Logistic regression was performed as used in other work [30], coefficients were fit separately for each session for each animal. In brief, we modeled choices of the mice on a trial to trial basis to understand its policy. The regressors used for the choice logistic regression were stimulus, previous choice, previous outcome and, the interaction term of choice and outcome. The added constant term represented the bias. The stimulus regressor was crafted taking the logarithm of tone frequencies in Hz and then normalizing it.

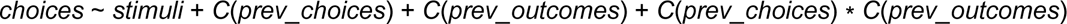

#### Psychometric fits and artificial classifiers

Psychometric curves were fitted to further characterize behaviour and performance during generalization tests. Curves corresponded to the probability of choosing category B (high frequencies) as a function of normalized log-distance of tone frequencies. The functional form corresponded to the error function with three adjustable parameters (bias, slope and lapse rate) obtained with maximum likelihood estimation [31]. Behavioural decision data and psychometric fits were also compared to resulting decision gradient curves of machine learning classifiers. A list of classifier algorithms as provided by library Sklearn (built-in-functions), was trained on a single session of the first phase of the experiment (discrimination stage) and then tested on a single session of the first generalization test, as completed by the mice subjects.

#### Single cell analysis

To define cells as modulated by stimulus or choice event we considered 1 s before and 3 s after and aligned to the event of interest. We used a parameter-free method called Zenith of Event-based Time- locked Anomalies (ZETA) [32]. It is agnostic to the shape of the modulation pattern elicited and makes no assumptions on the temporal modulations distribution. It computes if a neural activity pattern elicited by an event is statistically significant by checking if it could have been observed by chance. This pattern is compared to the likelihood of observing it in the null distribution built by running multiple bootstraps at jittered event-onset times randomly selected as pseudo-events. Then, if the direct percentile position of the peak anomalies is below 5 percent (p < 0.05, two-tailed) then the cell was labeled as modulated.

#### Response profile analysis

We concatenated the average responses (peri-event time histograms) of each neuron for each trial type (according to category, choice and outcome) to generate a vector. Next, dimensionality reduction was applied through Principal component analysis (PCA) to reduce complexity while preserving 90% of the variance. Then, we performed a t-SNE, short for t-Distributed Stochastic Neighbor Embedding, on the reduced data to visualize this high-dimensional data into two dimensions using euclidean distance as the metric. Silhouette values [33] were computed for different k-cluster values to evaluate presence of subpopulations with stereotypical responses in this reduced two-dimensional response profile space.

#### Population decoding analysis

We aligned neuron activities to the stimulus or choice onsets and trained individual decoders (linear support vector machines, linear-SVM, scikit-learn Python library) for every time step in a window (in trial time) that ranged from -0.8 s to 2.5 s for stimulus and -1.5 s to 2.5 s for choice around the alignment event. To build the training and test datasets, we used the pseudopopulation method, as described elsewhere [13,17,18]. In brief, for each subject and session, we divided data into trials of each of the four value trial type combinations. Next, neural activity was binned into 100 ms bins taking the average activity for each neuron recorded across the window. For cross-validation, we randomly divided data into two non-overlapping groups of trials, for training and testing the classifiers (a split of 70% trials was randomly selected for training, leaving the remaining 30% for testing performance). To avoid confounding factors during decoding of each variable, trials corresponding to each trial type combination were split separately [18]. Then, for each combination and session, we randomly sampled population vectors from the corresponding trials in the training or testing set. To balance the number of different trial types, trials were resampled with replacement to ensure a feature matrix of dimensions 2n by n for each combination, with n being the number of sampled neurons for that session. To extend this to build a pseudo-population dataset integrating different sessions and subjects, we performed the same procedure and then concatenated for each different session. Since decoding accuracy levels depend on the number of neurons used as features, to compare between phases of the experiment, the same number of neurons was sub-sampled for each session [17,34]. Hence, the final matrix would have total dimensions of 8N by N with N being the total number of neurons. Finally, neuron activity values both in training and testing matrices were z-scored across trials (for each neuron) according to the mean and standard deviation obtained from the training set. We repeated the whole analysis process multiple times to get a cross-validated estimate as the average (typically 100 times, if not reported otherwise). To compute significance at each timepoint we built a null distribution by performing the same decoding procedure but with permuted labels (500 times). Then, the p-values were computed by direct percentile position of the observed decoding accuracy with respect to the null distribution taking the maximum of the corresponding percentiles across timepoints to correct for multiple comparisons. Unless stated otherwise, the decoding analysis was performed using this pseudo-population method, aggregating recording sessions from the same stage and across mice subjects and days.

#### Video choice decoders

To extract animal behavioral features we used the DeepLabCut software to track five points of the animal: tongue, left port, right port, nose and back. Around 6% of the sessions were annotated manually, after automatic selection of representative frames by the in-built clustering algorithm of the preprocessing pipeline in DeepLabCut. The output traces were used to extract several variables of interest. Licking side and responses for each trial were extracted from the tongue and two port markers, including anticipatory licking when no ports were in reach. We also extracted general unease of the animal using the back marker as a coordinate to generate a region of interest (ROI). Next, this ROI was used to compute the normalized difference of the matrix norm of the video array frame to frame. In this way, by analyzing significant peaks in this resulting process, frames were labeled as with movement or not. These were later used in the behavioral decoders predicting choice as regressors.

#### Difference across decoding accuracy traces

To determine whether two decoding accuracy traces belonging to different stages were actually different, we first computed the absolute difference (delta) by subtracting the mean traces from one another at each timepoint. For each timepoint, we computed significance with respect to a null distribution of differences in direct percentile position. This null distribution was built by taking the cross-validation run traces of the stages of the pair of interest and shuffling them before splitting them generating two pseudo-stage groups to compute the delta difference in the same way as for the real grouped data. This entire process was repeated 500 times. Finally, we tackle the multiple comparisons problem across timepoints with Bonferroni correction.

#### Generalization decoding analysis

To study the geometry of the representations of trial type neural states, we made use of cross-condition generalization performance, as described in Bernardi <u>et al</u> [13] and applied it to every given trial timepoint to study the dynamics within the trial. Additionally, we preserved each split, i.e. each cross-pair of variables, to further understand the geometrical arrangement. In brief, we split the training and testing sets to decode a specific variable (main variable), according to the trial labels given by another one (split variable) such that testing data came from different conditions not used in training. For example, to test generalized encoding of choice, the decoder was trained to discriminate right (R) and left (L) choice in rewarded (Rw) trials and its performance was tested on discrimination of right (R) and left (L) choice in non-rewarded (NRw) trials and vice versa. The mean of these cross-pairs would then correspond to decoding choice with outcome split.

#### Rectangularity index

We computed the index to quantify how rectangular was the geometry arrangement of trial types on neural state space projected on the decoding plane of choice and outcome by the product of two terms. One accounting for the lengths of the segments between the means of the trial type states (sides of the rectangle) and another one to account for the angles between adjacent segments. The index would then range from 0 to 1, with 1 corresponding to a perfect rectangle with equal opposite sides by pairs and angles between adjacent sides being 90 degrees. The rectangularity index was computed following this expression:

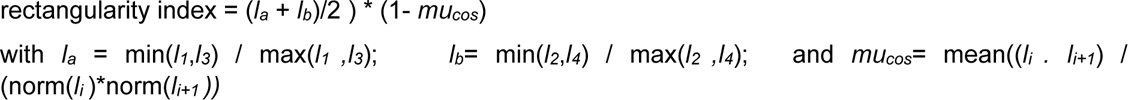

#### Simulation neural state space

To support our findings in the decoding and geometry analysis, we analyzed the geometry arrangements of simulations constrained by their decoding properties. We generated ∼90000 simulations of neural state space arrangements representing 4 trial types in an embedding space of three dimensions. The generative parameters were: rank of the matrix of state space vectors (to each trial type state mean) determining dimensionality of the arrangement, relative spread (sigma) of trial states around each trial type mean characterizing state noise, entropy of distances between trial state means to the centroid and, entropy of noise of trial type states. Generating a simulation given a specific dimensionality did not enforce a particular geometrical arrangement since independent vectors in the matrix were generated at random and the distances between them did not imply specific symmetries or angles between them. In other words, we did not enforce particular geometries.

#### Binomial regression model

We fitted a series of nested binomial generalized linear models (GLM) of the form:

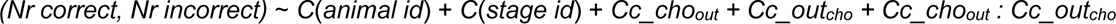

with a logit link to predict the fraction of correct choices (Correct and Incorrect counts) across 6 animals in 4 stages (St_id), using standardized neural activity parameters (Cc_cho, Cc_out) and their interaction.The model was implemented in Python with statsmodels (covariance type HC3) and Patsy libraries with GLM built-in function with Binomial family, with the response variable specified as a two-column DataFrame of successes and failures. The dataset comprised 22 observations (one per animal-stage tuple).

- Subject effect: To test whether subject id was relevant in explaining variance observed in the data, we computed the Intraclass Correlation Coefficient (ICC) which measures the proportion of total variance in the response variable that is attributable to differences between subjects. It quantifies how much of the variability in the fraction of correct choices (Fr_corr) is due to between-animal differences versus within-animal variability across stages. The between-animal variance for fraction of correct choices was 0.0014, and the within-animal variance was 0.0060, yielding an ICC of 0.19, indicating moderate clustering. However, sample sizes (nr datapoints < 30) in mixed models can lead to overfitting and poor generalizability and estimating a stable variance for each subject can be unreliable [35]. An ICC of 0.19 is below common thresholds (< 0.3) where random effects are typically recommended to account for substantial clustering, indicating that fixed-effects models can be used as an alternative and may suffice unless clustering is a primary focus [36,37]. Hence, the choice for the model included fixed effects to model subject effect balancing fit, parsimony and generalizability, focusing on the primary predictors to test our hypothesis (Cc_choout + Cc_outcho and their interaction) rather than animal-specific variation. A mixed-effects model could be considered if the ICC exceeds 0.3, but the current fixed-effects model is appropriate given the sample size and modest clustering.
- Interaction term: The interaction term between Cc_choout + Cc_outcho was introduced in the model explicitly as to test our hypothesis about its relevance, in comparison to each of the cross decoding accuracies alone, relating to different geometry arrangements. To test whether this actually constituted a significant advantage, we used the likelihood ratio test (LRT) for nested models, comparing each model with the interaction term against the corresponding reduced model without it. Other models without interaction term were also included in the modelling process and selected according to the same parameters of explanatory power.
- Multicollinearity between regressors: We examined predictor multicollinearity by computing Variance inflation factors (VIFs) for each regressor. VIF measures for each regressor how much of the variance of its estimated regression coefficient is increased due to multicollinearity with other predictors. Regressors with VIFs above 5 were considered collinear, following previous work [38,39].
- Standardization of regressors: We used standardized regressors for the neural predictors (Cc_choout + Cc_outcho and their interaction) so as to obtain the effect in normal units. This choice addresses collinearity between regressors (that have a small range in raw values), drastically reducing VIFs. It also reduces the risk of numerical instability during the estimation process and facilitates interpretation in terms of standard deviation units. It aids comparability across subjects and reduces scale-related bias in the interaction term. However, as a double check we also ran the final selected model for the raw regressors obtaining the same qualitative results.
- Model validation: Goodness of fit was assessed with the Pseudo R-squared for generalized linear models (GLMs), analogous to R-squared in linear models, as given by in-built function GLM() in Statsmodels (Cox-Snell pseudo R square), as well as, log likelihood and residual deviance. Explanatory power was tested in relation to the complexity of the model to favor parsimonious models, by using Bayesian Information Criterion that derives from log likelihood but also accounts for number of parameters in the model as well as number of data points [40]. To further assess how appropriate the complexity of the model was, we performed the deviance-based test for dispersion by dividing the residual deviance by the residual degrees of freedom [41] to estimate the dispersion. This statistic is approximately chi-squared distributed with df = n (number of datapoints) − p (number of parameters) under the null hypothesis of no over- or under dispersion. A large deviance relative to the degrees of freedom suggests poor fit. If it is > 1, the model shows overdispersion, meaning it expects less variability than what is actually present in the data (overly simple model). On the other hand, < 1 means under dispersion where the data presents less variability than expected by the model (overly complex model). Finally, to assess potential overfitting in the binomial regression model, we computed the leave-one-out cross-validation (LOOCV) mean squared error (MSE). LOOCV involves fitting the model n times, each time excluding one observation, using the remaining n−1 observations as the training set, and predicting the held-out observation. The LOOCV MSE is the average of the squared prediction errors across all n iterations, providing an estimate of the model’s performance on unseen data. Typically, LOOCV MSE is expected to be higher than in sample MSE. By comparing the gap between both (optimism) and comparing it to the LOOCV MSE variability (standard deviation) we evaluated if there was relevant overfitting [42].
- Residual diagnostics and influence: To assess the influence and potential outliers in the binomial regression model, we computed deviance residuals, leverage, and influence for each datapoint using in-built functions of the GLM Influence object from the Statsmodels library in Python. Deviance residuals measure the contribution of each datapoint to the model’s overall deviance, which quantifies the difference between the fitted model and a perfect (saturated) model. For a single observation, the deviance residual is defined as the signed square root of the observation’s contribution to the total deviance, adjusted to reflect whether the observed value is above or below the predicted value [41]. Leverage measures how much a datapoint’s predictor values (independent variables, i.e. regressors) deviate from the mean of those predictors. High leverage points are outliers in the predictor space, meaning they have unusual combinations of predictor values. Points with high leverage can disproportionately influence the model’s fit, even if their response values are not extreme. In this study, we considered data points with leverage above the threshold 2*p/n, with p being the number of parameters of the model and n the number of datapoints, as extreme in the predictor space and prompting further analysis of deviance and influence [43]. Lastly, to measure the influence of each datapoint we computed Cook’s distance that combines leverage and residuals to assess how much the fitted model would change if a particular point were removed. In binomial regression, Cook’s distance is calculated as a function of the standardized residual and leverage, with higher values indicating greater influence. We used a distribution-based cutoff using the median of a chi-squared distribution with as many degrees of freedom as the number of parameters in the model, following Kim et al. [44]. By combining these parameters, a datapoint was considered an outlier when both surpassing the influence threshold (Cook’s distance) and leverage. Additionally, by plotting deviance residuals against actual fitted values, we analyzed evidence for non-linearity, heteroscedasticity, overdispersion or any other patterns of the model [41,43,45].

Among all fitted models, those with high VIFs were discarded. After this first constraint, the final model was selected upon minimizing BIC.

For the model finally selected, the formula was:

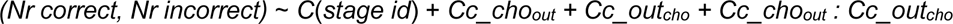

VIFs ranged from 1.11 to 4.67, all below the threshold of 5, indicating no severe collinearity. A correlation heatmap of regressors confirmed this (Fig. S6), with pairwise correlations between continous regressors being modest: Cc_choout and Cc_outcho at 0.41, Cc_outcho and Cc_choout:Cc_outcho at -0.14, and Cc_choout and Cc_choout:Cc_outcho at 0.06. Correlations involving St_id dummies were similarly low, with no absolute values exceeding 0.4. These results corroborate the VIFs, demonstrating that multicollinearity does not compromise coefficient estimates or model stability significantly. The model achieved a deviance of 21.18, compared to a null model deviance of 80.08, explaining 58.9 units of deviance (73.6% reduction). A likelihood ratio test (LRT) confirmed this improvement (χ² = 58.9, df = 6, p < 0.001), and also the advantage of including the interaction term by testing LRT respect to a reduced model without the interaction term (χ² = 35.81 - 21.18 = 14.63, df = 1, p < 0.001). The Pseudo R-squared (CS) was 0.9313, indicating strong explanatory power. The log-likelihood was -63.80, and the BIC was 149.24. Coefficients (Fig. 5H) showed significant effects for Cc_choout (β = -0.1556, p = 0.001) and the interaction Cc_choout : Cc_outcho (β = 0.1754, p < 0.001), with stage effects varying in significance (e.g., C(St_id)[T.gentest_1], β = -0.3685, p = 0.006). Standardized residuals ranged from -2.54 to 2.93, and observation datapoint with index 14 qualified as outlier with deviance residual of 2.93 (> 2) and leverage 0.58 (> 2p/n) with a resulting Cook distance of 1.69. A plot of residuals versus fitted values showed no systematic pattern, supporting model adequacy. Same model was also fitted without the outlier observing no qualitative changes on significant neural parameter regressors or sign of coefficients (Deviance explained: 58.66, BIC: 143.46, Cc_choout (β = -0.1556, p = 0.001) and the interaction Cc_choout : Cc_outcho (β = 0.1755, p < 0.001).

The in-sample MSE between fitted probabilities and observed fractions (Fr_corr, range: 0.637 to 0.918) was 0.0015, while the LOOCV MSE was 0.0056, yielding an mse gap (optimism) of 0.0041.To assess overfitting in the binomial regression model, we employed a bootstrap-based test to estimate the optimism (MSE gap) distribution, as described by Hastie et al. [42]. For 1000 bootstrap samples, the model was trained on resampled data, and the in-sample MSE was compared to the out-of-sample MSE on the original data to compute the optimism. The observed MSE gap was then evaluated against the bootstrap 95% confidence interval (-0.000126 to 0.008559). As a double-check, a reduced model with fewer predictors was also tested (after removing St_id regressors), yielding an observed MSE gap of 0.002294 (64.89% relative increase) and a 95% confidence interval of (-0.000361 to 0.005072). Both models exhibited qualitatively similar results, with the same coefficients identified as statistically significant, confirming consistency in the key neurally derived predictors.

The dispersion parameter was 1.41 following the Pearson chi-squared statistic (21.2) relative to 15 residual degrees of freedom (χ²/df = 1.41) suggested no severe overdispersion. Moreover, its deviance from 1 did not reach significance under the chi square distribution with respect to number of data points and parameters, concluding an acceptable generalization performance [41].

### Software and Statistical Tests

For all image processing, automation, data analysis and statistics we used the Python programing environment with the help of packages like Datajoint, CaImAn, DeepLabCut, DeepCadRT, Patsy, scipy, statsmodels and Sklearn, among others. Summary data are reported as mean ± s.e.m. (standard error of the mean) unless otherwise specified.

Box plots indicate median (center), 25th and 75th percentile (box) and most extreme data points (whiskers) that were not considered outliers (points for which the distance from the box exceeds 1.5 times the length of the box). Violin plots indicate mean (center) and minimum and maximum values (whiskers) of the distribution.

Statistical tests were performed at the group level with parametric testing for behavioral variables unless specified. Normality of data distributions was determined by the Shapiro-Wilk test via the built-in-function available in the spicy Python library, applying non-parametric alternatives when violated. Threshold for statistical significance was defined as 0.05. Bonferroni correction was applied when doing multiple comparisons. For the neural activity analysis, p-values were computed using a null distribution constructed from shuffling labels and using a direct percentile position.

Spearman rank was used when testing correlation for two variables unless otherwise specified.

No power analysis was run to determine sample size a priori. The sample sizes chosen are similar to those used in previous publications. We excluded three animals because of weak signals of neuronal activity, due to insufficient labeling and/or GRIN lens misplacement. We had to terminate the experiments for 4 mice to be in accordance with the animal welfare regulations because they presented with unbiasable response policy despite extended training or did not progress in the staging in the available experimental time. Only mice that completed all stages of the experiment were used for final analysis (n=6).

## Acknowledgements

We would like to thank Thomas Akam, Jorrit Montjin, Lorenzo Posani and Andrew Macaskill for feedback about data analysis and methods. Additionally, Simon Mussal, Pietr Goltstein, Adrian Roggenbach and Shuting Han for fruitful discussions about study design and animal training. Finally, we would like to acknowledge the help from Simone Holler-Rickauer for the preparation of the histology slides.

## Author Contributions

L.S.V. and B.F.G. conceptualized the study. L.S.V. carried out all in vivo imaging experiments, preprocessing of data and data analysis. L.S.V. build and automated the behavior setup. V.M. provided conceptual feedback for the neural data analysis and reviewed the final manuscript. R.B. helped with the animal preparation and histology and reviewed the final manuscript. I.C provided feedback on the neural data and statistical analysis, reviewed and edited the final manuscript. L.S.V. wrote the original draft and L.S.V. and B.F.G wrote, reviewed and edited the final manuscript. Funding acquisition and supervision were done by B.F.G.

## Data and code availability

Data is available upon request. Code is available at github.com/lsainzvillalba/hpc_cat_2025_paper.

