## supplementary figures for "Category learning disentangles representation of trial events in hippocampus CA1"

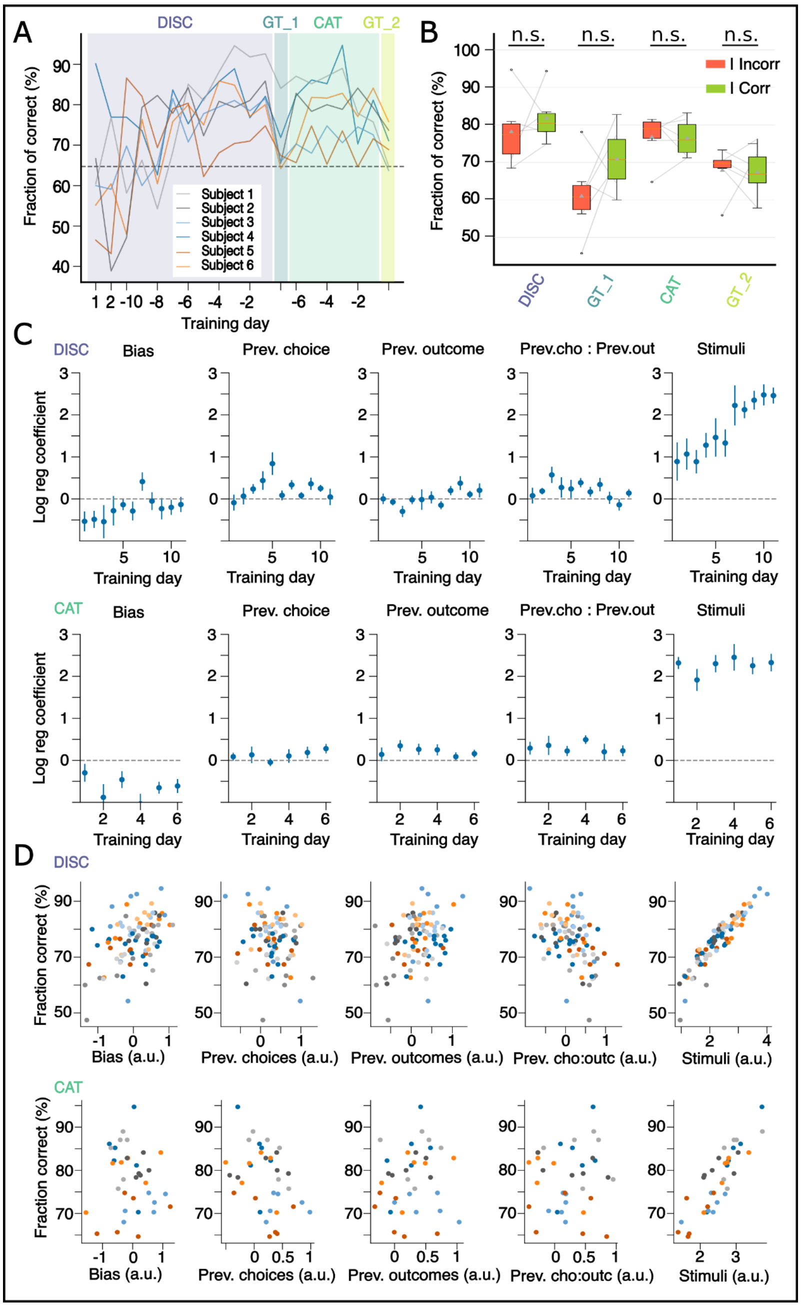

**Supplementary figure 1**: Overview of training and factors determining behavioral choices. (A) Training curves for each subject n=6, with shaded color for stages of paradigm. (B) Statistical comparison of fraction of correct choices after incorrect or correct trials across stages showing no significant differences. This supports choice strategy not strongly depending on the previous outcome of first order (double-sided, paired t-test). (C) Trend in logistic regression coefficients predicting choice on a trial to trial basis for discrimination stage (top) and categorization stage (bottom). The only regressor showing significant trend corresponded to the one weighing stimulus information in the discrimination stage (n=6, bias: Q =11.909, p=0.291, previous choices: Q=17.667, p=0.061, previous outcomes: Q=13.909, p = 0.177, previous choice-outcomes: Q= 13.152, p= 0.215, stimuli: Q=29.667, p<0.001, Friedman Rank Sum test). For the categorization stage, none exhibited significant changes (n=6 bias Q=6.857 p=0.231, previous choices Q=1.429, p=0.921, previous outcomes Q=2.667, p = 0.751, prev choice-outcomes Q= 2.381, p= 0.794, stimuli Q=3.810, p=0.577, Friedman Rank Sum test). However, for the stimulus regressor, the coefficient was stable at a significant high value. (D) Correlation test scatter plots for each logistic regression regressor and fraction of correct choices for discrimination stage (top) and categorization stage (bottom). Color code corresponds to animal id. Spearman rank correlation coefficient was computed for all datapoints for all subjects. If significant, a control test (t-test for independent samples) was performed across subjects on the rho coefficient computed separately for each subject and every regressor to avoid aggregation bias (also called Simpson's paradox) (discrimination: n=6, bias rho: 0.037 p=0.769, previous choices rho: -0.101 p=0.419, previous outcomes rho: 0.342 p=0.005, t-test p=0.008, previous choice-outcomes rho: -0.261 p=0.035, t-test p=0.927, stimuli rho: 0.974 p<0.001, t-test p<0.001 ) (categorization: n= 6, bias rho: -0.045 p = 0.794, previous choices rho: -0.269 p=0.113, previous outcomes rho: 0.010 p=0.956, previous choice-outcomes rho: -0.112 p=0.514, stimuli rho: 0.868 p<0.001, t-test p<0.001)

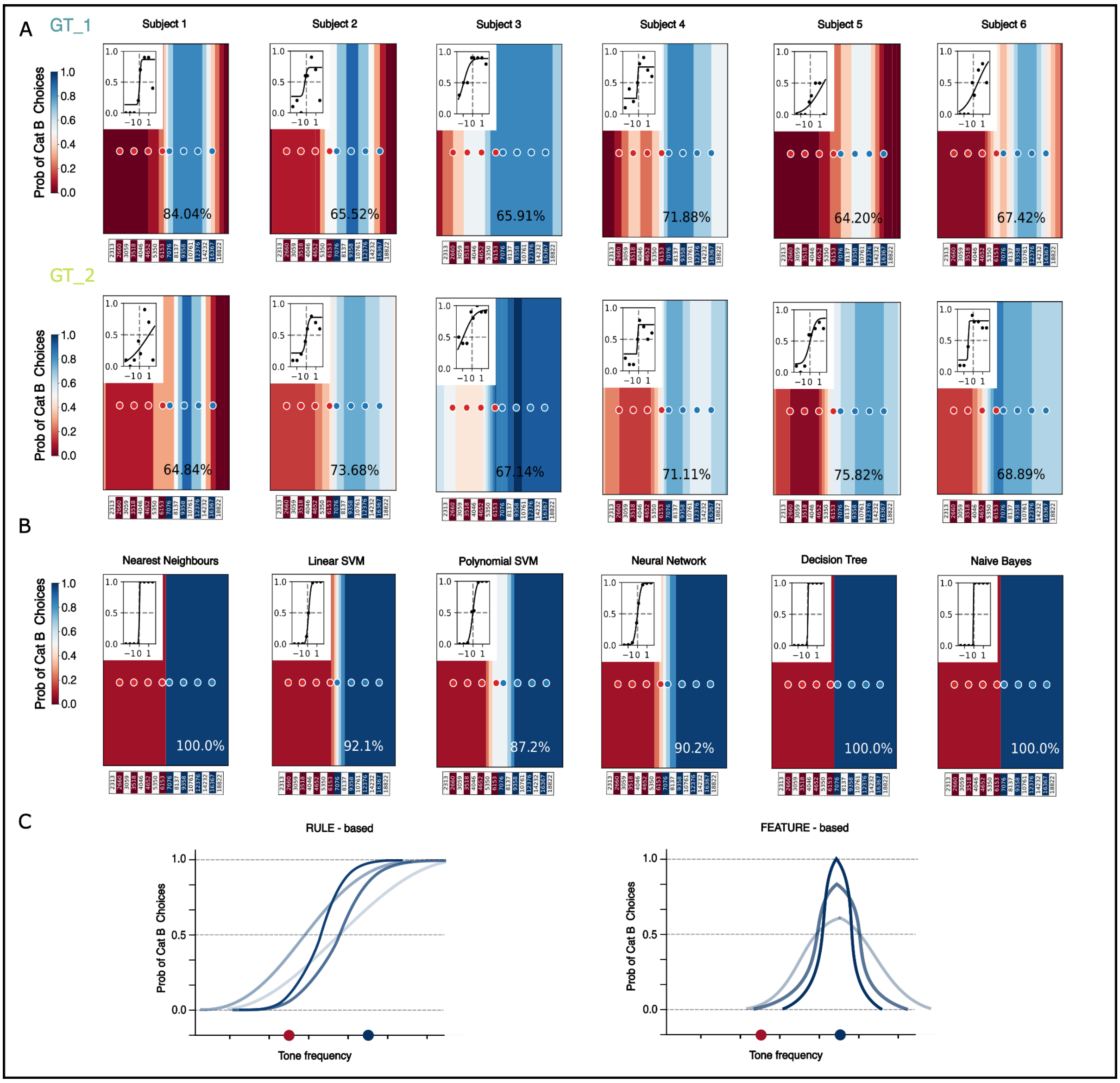


**Supplementary figure 2**: Psychometric fits and decision-function plots for all subjects and examples of artificial classifiers. (A) Each column displays decision-function heatmap plots with corresponding psychometric fit (upper left corner) for each subject in either generalization test 1 (GT_1, top) or generalization test 2 (GT_2, bottom). Some subjects develop biased responses during test sessions like Subject 3 or Subject 6, other subjects improved from generalization test 1 to 2. (B) Artificial classifier decision-function plots trained in discrimination phase stimuli and tested for generalization using the same stimuli set as in generalization tests applied to mice subjects. All of the examples, without further tuning, saturate at the edges of the stimuli space, far from the boundary. (C) Illustration schematic for different decision curves assuming rule-based generalization (left) vs feature-based generalization (right). For rule-based generalization, stimuli far from the boundary exhibit a saturation in choice (probability choice distribution is skewed to one of the options, reflecting that these stimuli are easier to classify). In feature-based generalization, the likelihood of category membership, as given by choice probability, reflects the fact that the ability to generalize is tied to the similarity (in feature space) the new example has with previous experienced examples as in a density fitting algorithm. Different curves depict different possible kernels or different widths centered and fitted around experienced specific features. At the edges of the stimuli feature space (corresponding to extrapolation stimuli), rule-based and feature-based curves diverge. Colored circles in the x axis represent experienced features color-coded by their category.

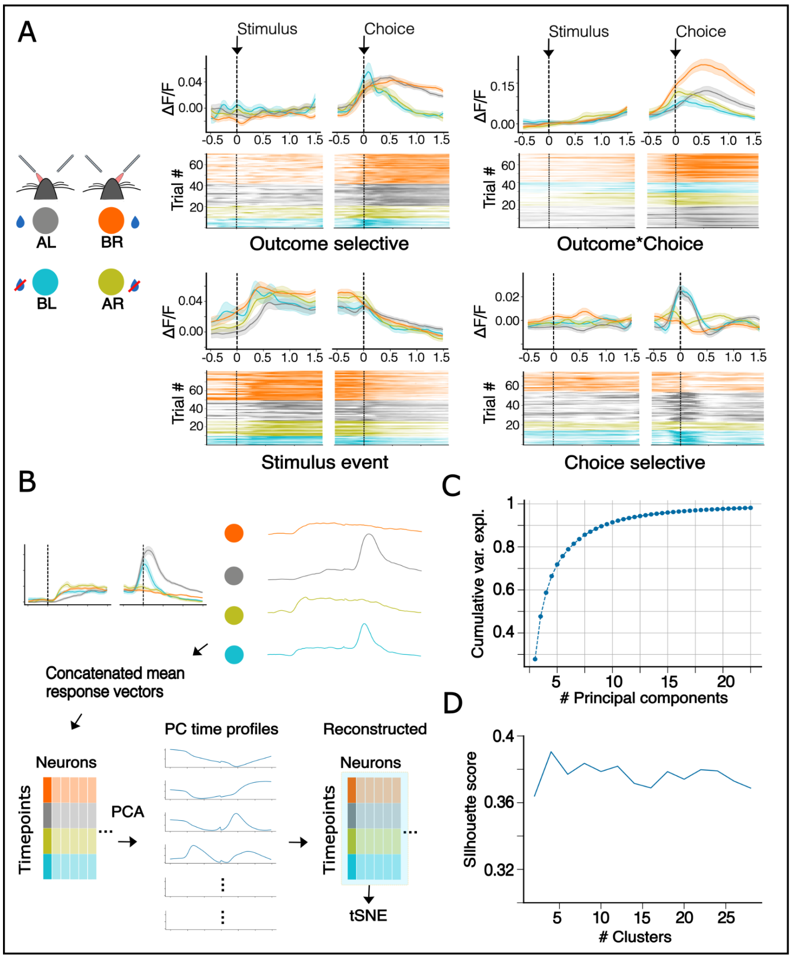


**Supplementary figure 3**: Single neuron response example profiles and t-SNE analysis. (A) Single unit example activity peri-event time histograms (PETHs) showing different response profiles for events in trial (like for stimulus presentation), or selective to certain value variables (choice or outcome) and even selective to a combination of both. (B) Processing method for t-SNE map. First mean PETH responses for each trial type are concatenated in one single vector for each neuron to create a matrix of timepoints x neurons. Principal component analysis (PCA) is performed to capture most relevant response time profiles. The matrix is then reconstructed on the basis of PCA, truncated to capture at least 95 % variance. We then obtain the t-SNE map running it on this reconstructed matrix. (C) Cumulative variance explained as a function of the number of principal components. 14 principal components capture at least 95% of variance and were used for reconstruction of the matrix. (D) Silhouette score computed for different numbers of clusters, showing no significant clustering (silhouette score <0.5) for any number of clusters tested (2 to 28) (p>0.05 , permutation test).

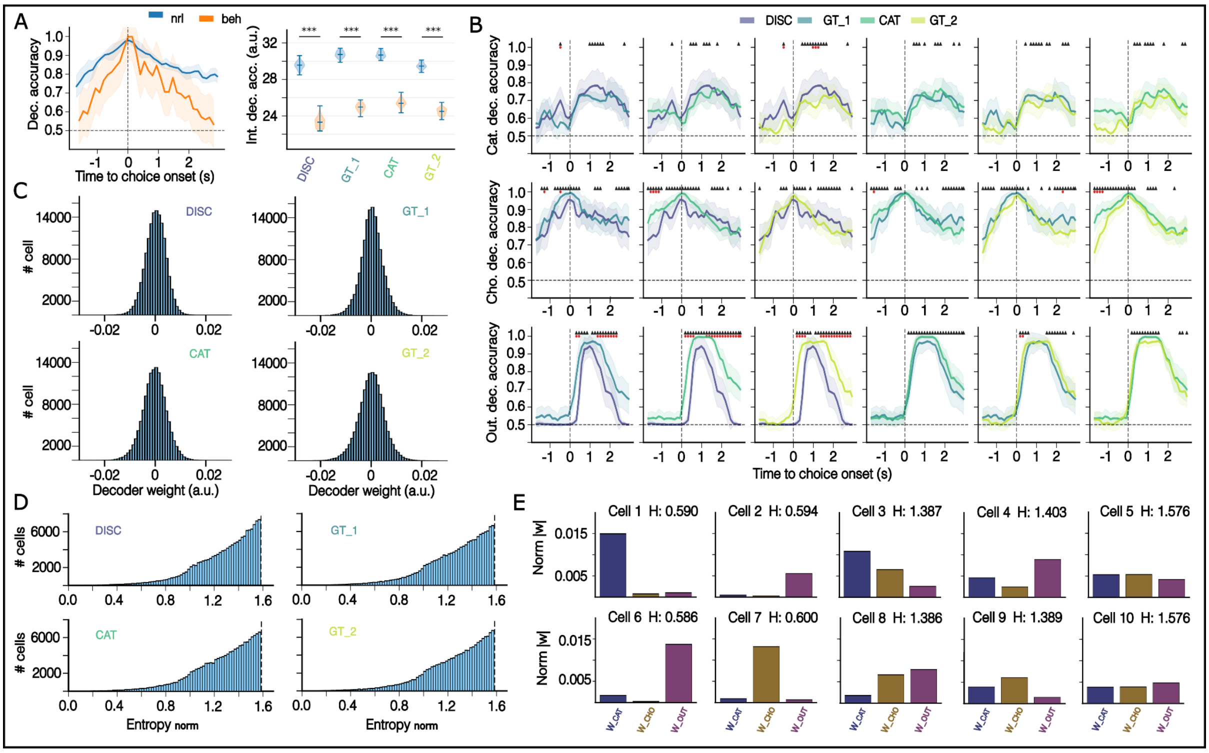

**Supplementary figure 4**: Decoding comparison and decoder weights. (A) (Left) Decoding in trial time for choice using neural activity (blue) and behavioral footage (orange) decoders. (Right) Comparison of integrated decoding accuracy for neural and behavioral decoders for choice across stages showing neural decoders persistently overperform behavioral ones (Mann-Whitney test, DISC: $p<0.001$, GT_1: $p<0.001$, CAT: $p<0.001$, GT_2: $p<0.001$). (B) Pair-wise stage comparison of decoding traces in trial time for category, choice and outcome variables. Black triangles represent timepoints with significant difference between the two stage traces (percentile on null distribution by bootstrap permutation test) for significant decoding timepoints. Red dots represent timepoints with relevant effect size: > 10 % decoding difference among the significant timepoints. Most prominent differences occur for outcome decoding, particularly between discrimination and the rest of stages, in the post-choice window with high decoding accuracy increasing in time across stages. (C) Decoder weight distributions for outcome decoding in the different stages for all cross validation runs. All decoder weight distributions, including corresponding ones for choice and category followed a Gaussian distribution (Anderson-Darling test, $p< 0.05$) (D) Decoder weight entropy distribution across stages. For each cell, entropy is computed according to their decoder weights for category, choice and outcome as a normalized vector. Distributions show few cells with low entropy. Maximum entropy corresponds to equal weights among decoders $H_{max} = \log_2(3) = 1{,}585$. (E) Example cell decoder weight profiles with corresponding computed entropy. Low entropy corresponds to more selective profiles.

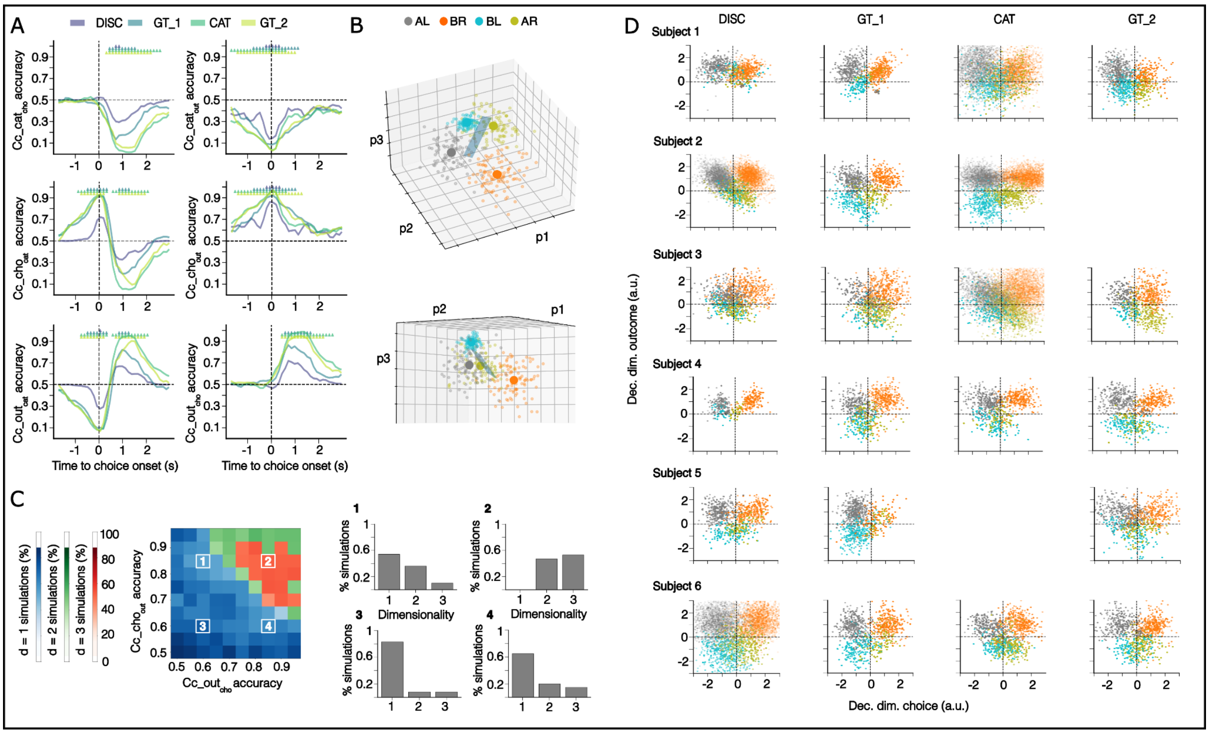


**Supplementary figure 5**: Cross condition decoding and neural state simulations. (A) Cross condition decoding for all variables by split pairs for all stages DISC: discrimination, GT_1: generalization test 1, CAT: categorization , GT_2: generalization test 2. Colored triangles indicate timepoints with significant decoding accuracy with respect to a null distribution (percentile on permuted labels) . Mirror-shaped traces by pairs of combinations show that when decoding for category or with category split, the decoder predicted opposite labels capturing choice or outcome information in training. (B) Example of simulation of neural state space with dimensionality d=2 an added gaussian noise that has choice and outcome as represented axis of the arrangement. The decoding plane for outcome (top) and for category (bottom). This does not preclude getting a high and above-chance decoding accuracy for category. (C) (Left) 2-d histogram of retrieved simulations when constrained to values of cross-condition decoding accuracies of choice and outcome color-coded by most common dimensionality for each combination of Cc_cho$_{out}$ and Cc_out$_{cho}$. (Right) Histograms (in percentage) of dimensionality corresponding to the numbered combinations of Cc_cho$_{out}$ and Cc_out$_{cho}$ in the 2-d histogram. For high accuracy values of Cc_cho$_{out}$ and Cc_out$_{cho}$ > 0.7, retrieved simulations have predominantly d=2 or d=3 with few or none 1-d simulations (panel # 2). (D) Neural space representation of trial activity color-coded by trial type as in Fig. 6E computed for each subject and projected on the axis of decoding dimensions for choice (x-axis) and outcome (y-axis). These decoding vectors are built as a coordinate basis from the weights assigned by the SVM classifier to each neuron serving as a feature. Some subjects present a clear progression on factorizing choice and outcome (Subject 1) and some others are more noisy with no full factorization by the last stage (Subject 5) or start already in an increased factorized state and seemingly remain stable (Subject 6). Blank plots correspond to missing sessions due to software imaging incidents (ScanImage error on recording).

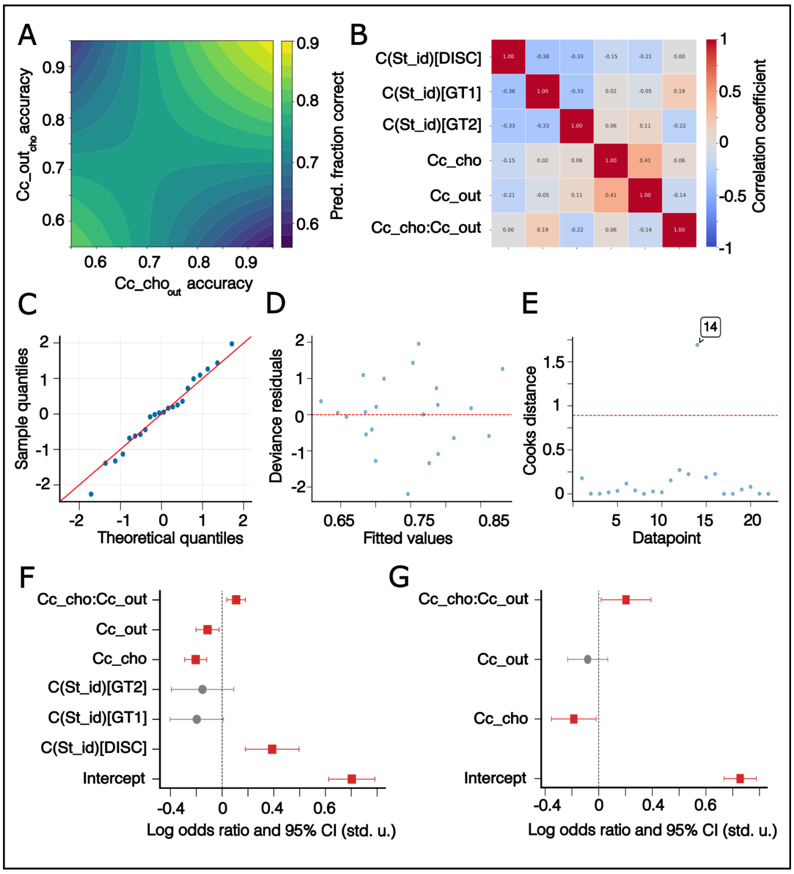


**Supplementary figure 6**: Binomial regression model results and validation. (A) Predicted fraction of correct heatmap to show interaction effect between Cc_cho$_{out}$ and Cc_out$_{cho}$ . Net positive effect starts to be relevant for accuracy values > 0.75, similar to as shown in simulations and aligned with theoretical interpretation of decodability of four balanced trial types corresponding to 2-d and 3d arrangements. (B) Correlation between regressors of the selected model. None of the pair-wise correlations is above 0.5, corresponding to variance inflation factors < 5, reflecting no major collinearity. (C) Q-Q plot between sample quantiles and theoretical quantiles for all datapoints. Residuals follow normal distribution (Shapiro-Wilk test). (D) Deviance residuals vs fitted values. No clear pattern of overdispersion is noted (no funnel pattern)  (E) Cooks distance for every datapoint. Datapoint number 14 is the only one surpassing the established threshold (see Methods) and hence labeled as outlier (F) Forest plot for selected model but fitted without the outlier. No qualitative changes occur as compared to the model fitting all datapoints. The regressor for Cc_out$_{cho}$ becomes significant but is similar in magnitude and with the same sign as in the full model. (G) Forest plot for reduced model, dropping the categorical regressors corresponding to stage id. Once again, there are no qualitative changes in neural parameter regressors with persistence of positive and significant contribution of interaction regressor.
